# Immunogenomic landscape of hematological malignancies

**DOI:** 10.1101/618918

**Authors:** Olli Dufva, Petri Pölönen, Oscar Brück, Mikko Keränen, Juha Mehtonen, Ashwini Kumar, Caroline Heckman, Sanna Siitonen, Kirsi Granberg, Suvi-Katri Leivonen, Leo Meriranta, Sirpa Leppä, Matti Nykter, Olli Lohi, Merja Heinäniemi, Satu Mustjoki

## Abstract

Understanding factors that shape the immune landscape across hematological malignancies is essential for immunotherapy development. Here, we integrated over 8,000 transcriptomes and over 1,000 samples with multilevel genomic data of hematological cancers to investigate how immunological features are linked to cancer subtypes, genetic and epigenetic alterations, and patient survival. Infiltration of cytotoxic immune cells was associated with distinct microenvironmental responses and driver alterations in different cancers, such as *TP53* in acute myeloid leukemia and *DTX1* in diffuse large B cell lymphoma. Epigenetic modification of *CIITA* regulating antigen presentation, cancer type-specific immune checkpoints such as VISTA in myeloid malignancies, and variation in cancer antigen expression further contributed to immune heterogeneity. Prognostic models highlighted the significance of immunological properties in predicting survival. Our study represents the most comprehensive effort to date to link immunology with cancer subtypes and genomics in hematological malignancies, providing a resource to guide future studies and immunotherapy development.

## INTRODUCTION

Immune checkpoint blockade therapies are revolutionizing cancer therapy in several tumor types, demonstrating that the immune system can be successfully harnessed for effective anti-cancer treatment (Ribas and Wolchok, 2018). In hematological malignancies, immune checkpoint inhibition has demonstrated efficacy in classical Hodgkin’s lymphoma (CHL) (Ansell et al., 2015), and adoptive chimeric antigen receptor (CAR) T cell therapy has been successful in several B cell malignancies (Maude et al., 2014; Schuster et al., 2017). Allogeneic hematopoietic stem cell transplantation (allo-HSCT) is also considered to rely on the immune system by inducing the graft-versus-leukemia effect (Casucci et al., 2013). It is typical for immune-based therapies that only some cancer types or a subset of patients within a cancer type achieve responses. Therefore, rational patient selection based on the immune milieu of each tumor type may be crucial to achieve optimal benefit from immunotherapies. However, in hematological cancers the immunological diversity and underlying mechanisms resulting in distinct immune landscapes are unclear.

The immune landscape of cancers comprises various elements influencing the anti-cancer immune response (Chen and Mellman, 2013). The composition of the immune infiltrate, importantly cytotoxic lymphocytes that mediate elimination of cancer cells, has been associated with favorable outcomes in several cancers (Fridman et al., 2012; Galon et al., 2006) and with immunotherapy responses (Tumeh et al., 2014; Van Allen et al., 2015). Furthermore, antigen presentation is essential for adaptive immune responses. Antigen presentation by cancer cells is commonly considered to occur in the context of human leukocyte antigen (HLA) class I molecules, and somatic mutations in the HLA genes are frequent immune evasion mechanisms in solid tumors (Garrido et al., 2010; Shukla et al., 2015). However, as the normal cellular counterpart for most hematological malignancies is closely related to antigen-presenting cells (APCs), hematopoietic cancer cells may present antigen also in the context of HLA class II (Bachireddy et al., 2015). HLA II expression has been linked to prognosis (Rimsza et al., 2004) and response to PD-1 blockade immunotherapy in lymphoma (Roemer et al., 2018), and loss of mismatched HLA (Vago et al., 2009) or transcriptional downregulation of HLA II (Christopher et al., 2018) has been associated with relapse after allo-HSCT in acute myeloid leukemia (AML), demonstrating the importance of HLA II in hematological malignancies.

In addition to antigen presentation, T cells require costimulatory signals for effective activation. The balance of activating and inhibitory signals from tumor cells and APCs regulates both T and NK cell responses, and inhibitory signals such as PD-L1 are proven therapy targets (Chen and Flies, 2013). Finally, the presence of cancer antigens is essential to initiate and maintain the adaptive immune response. Cancer antigens include neoantigens derived from somatically mutated genomic regions and cancer-germline antigens (CGAs) normally expressed only in immune-privileged germ cells (Simpson et al., 2005). Several hematological malignancies have been shown to harbor low neoantigen loads (Alexandrov et al., 2013). However, CGA expression across hematological malignancies has not been systematically analyzed, although studies within cancer types have been conducted (Atanackovic et al., 2007; Meklat et al., 2007).

Emerging evidence from solid tumors suggests that cancer cell-intrinsic genetic and epigenetic aberrations influence tumor immune landscapes (Wellenstein and de Visser, 2018). Recently, several studies have integrated the genetics of solid tumors with immunological properties by leveraging extensive genomic datasets (Charoentong et al., 2017; Gentles et al., 2015; Li et al., 2016; Rooney et al., 2015; Thorsson et al., 2018). In contrast, large-scale studies of genotypic-immunophenotypic connections in hematological malignancies have not been performed. However, it is likely that heterogeneity exists in immunological properties given the vast genetic and epigenetic heterogeneity in hematological malignancies such as AML (Cancer Genome Atlas Research Network, 2013; Figueroa et al., 2010).

Here, we perform a comprehensive immunogenomic analysis in hematological cancers, investigating cytotoxic immune infiltration, antigen presentation, immune cell costimulation, and cancer antigen expression patterns in relation to cancer subtypes and genomics. We utilize a resource of over 8,000 transcriptomes collected across 36 hematological malignancies and normal hematopoietic cells (Hemap, http://hemap.uta.fi), together with multi-omics datasets of AML and diffuse large B cell lymphoma (DLBCL). In addition to transcriptomics, we integrate somatic DNA alterations, DNA methylation, quantitative multiplex immunohistochemistry, and flow cytometry to comprehensively map immunological features and validate the robustness of the findings. We identify microenvironmental differences between immune-infiltrated and immune-excluded cancers and demonstrate how the genetic and epigenetic makeup is linked to immune infiltration and antigen presentation. This understanding has implications for the development of precision immune intervention strategies in hematological malignancies.

## RESULTS

### Assessment of cytotoxic lymphocyte infiltration across hematological malignancies

We used 8,472 samples from 36 hematological malignancies, with 629 healthy donor hematological cell populations and 530 cell lines as controls, to comprehensively analyze immunological properties in hematological cancer transcriptomes (Figures 1A and S1A and Table S1). To facilitate linking immunological features to molecular cancer subtypes, we visualized the data using t-Distributed Stochastic Neighbor Embedding (t-SNE) (van der Maaten and Hinton, 2008) and utilized unsupervised sample stratification using density-based assignment of clusters with distinct molecular profiles (Cheng, 1995; Mehtonen et al., 2019).

**Figure 1.**
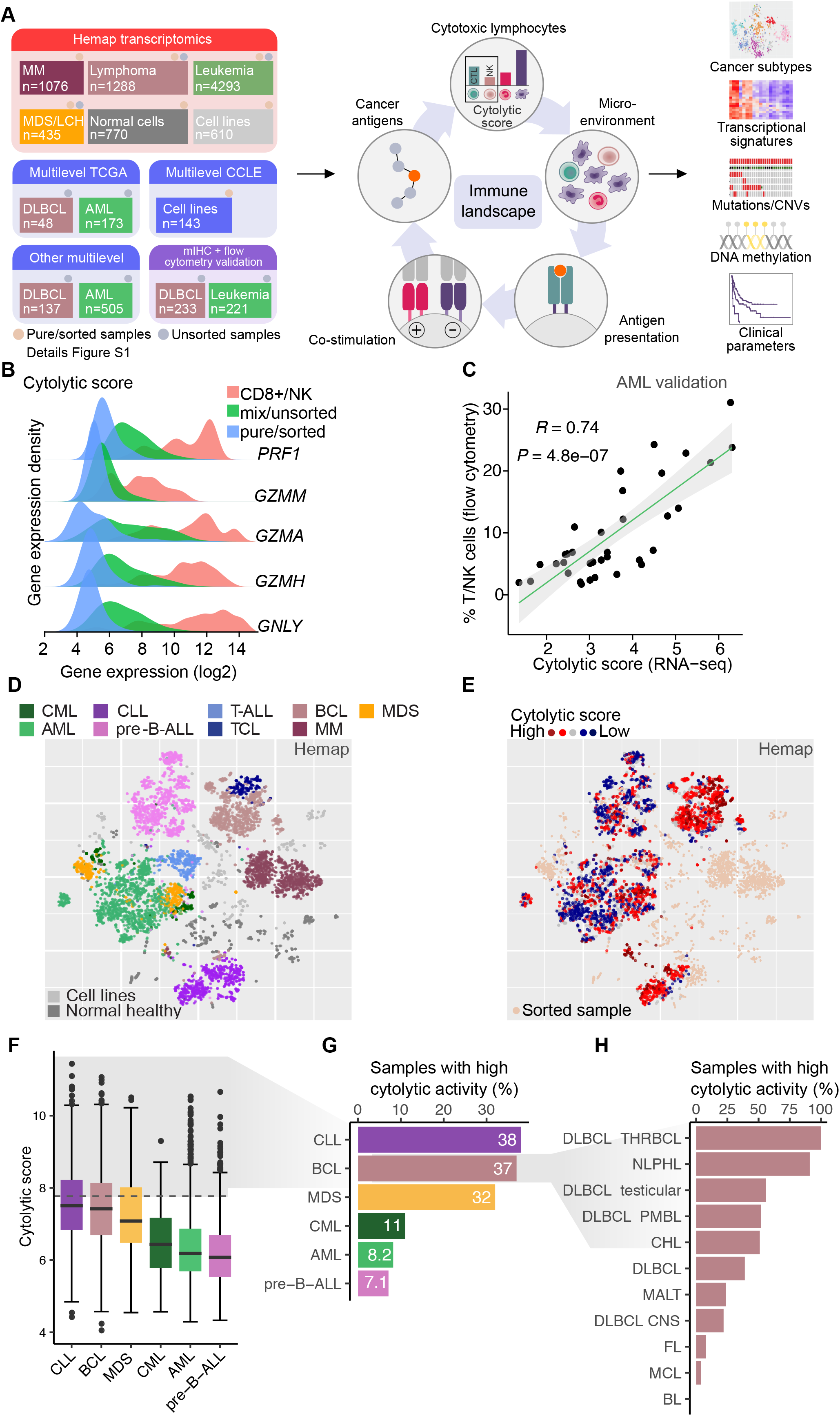
Identification of cytotoxic lymphocyte infiltration in hematological malignancies. **A**. Schematic overview of the study. Data from hematological malignancies and normal hematopoietic cells from Hemap and other sources were integrated to study associations of immune states to cancer subtypes, transcriptional, genetic and epigenetic properties, and clinical parameters. **B**. Distributions of expression levels (log2 expression) of genes included in cytolytic score for sorted cancer cells, unsorted tumor samples, and sorted CD8+ cytotoxic T lymphocytes and NK cells. **C**. Correlation (Spearman) of cytolytic score obtained from RNA-seq and combined T and NK cell fraction obtained by flow cytometry from 35 AML BM samples. **D**. Visualization of Hemap samples on a t-SNE map, with cancer types, cell lines, and normal cell populations colored. **E**. Cytolytic score colored on the Hemap t-SNE map. Red and blue color tones correspond to high and low scores, respectively. Sorted samples (score not calculated) are colored in beige. **F**. Cytolytic score across main cancer types in Hemap shown as box plots. Grey shading indicates samples with Z-score > 1. **G**. Percentages of samples with high cytolytic score (Z-score > 1) across main cancer types. **H**. Percentages of samples with high cytolytic score (Z-score > 1) shown as in G across Hemap BCL subtypes. See also Figure S1 and Table S1.

CD8+ cytotoxic T lymphocytes (CTLs) and natural killer (NK) cells are considered to be essential for effective anti-tumor immunity and responsiveness to immunotherapy (Joyce and Fearon, 2015; Morvan and Lanier, 2016; Tumeh et al., 2014; Van Allen et al., 2015). We first aimed to quantify the cytolytic immune infiltrate in the tumor microenvironment from bulk transcriptomes across hematological malignancies genes specifically expressed in CTLs and NK cells (Figure S1B). Based on the high specificity of the genes *GZMA, GZMH, GZMM, PRF1*, and *GNLY* to CTLs/NK cells compared to hematopoietic cancer cells and their essential role in cytolytic effector functions, we defined the geometric mean of these five genes as the cytolytic score reflecting CTL/NK abundance, (Figures 1B and S1C).

To validate our strategy of inferring cytotoxic lymphocyte abundance in hematological malignancies, we analyzed T and NK cell fractions using flow cytometry from diagnostic bone marrow (BM) aspirates of AML patients and performed paired RNA-seq from BM mononuclear cells. Cytolytic score correlated highly with the combined fraction of T and NK cells out of all BM cells, indicating good performance in leukemia samples (Spearman’s *R* = 0.74, *P* = 4.8 × 10^-7^, Figure 1C and S1D). Demonstrating utility also in lymphomas, cytolytic score correlated with the immunohistochemistry-based T cell content in a mucosa-associated lymphoid tissue (MALT) lymphoma dataset included in Hemap (Chng et al., 2009) (R = 0.68, *P* = 0.00013, Figure S1E). Built for hematological malignancies, cytolytic score also agreed well with previously reported methods of estimating immune cell subset abundance in solid tumors, including gene sets proposed by Bindea *et al.* (Bindea et al., 2013) and MCP-counter (Becht et al., 2016) (Figures S1F and S1G). Correlation to the deconvolution method CIBERSORT (Newman et al., 2015), designed to infer relative fractions of immune cell types rather than their abundance, was lower (Figure S1H). In conclusion, cytolytic score robustly estimates the abundance of CTLs and NK cells in bulk gene expression profiles of hematological malignancies, enabling its use for immunogenomic analyses.

Across transcriptomes of hematological malignancies, we observed the highest cytolytic score in chronic lymphocytic leukemia (CLL) and B cell lymphomas (BCL) (Figures 1D-F and Table S1). In contrast, acute leukemias and chronic myeloid leukemia (CML) were characterized by lower cytolytic scores. Importantly, we observed substantial variation in cytolytic infiltrate within cancer types, with most cancer types, including acute leukemias, harboring a subset of samples with high cytolytic score (Z-score > 1 across all cancers) (Figure 1G). Within BCL, we observed the highest levels in T cell/histiocyte-rich B cell lymphoma (THRBCL) and CHL, and the lowest in Burkitt’s lymphoma (BL) (Figure 1H), consistent with known characteristics of the immune infiltrate across BCL subtypes (Scott and Gascoyne, 2014). Activated B cell-like (ABC) DLBCL showed higher cytolytic score compared to the germinal center B cell-like (GCB) subtype. T cell lymphoma (TCL) subtypes had high cytolytic score, likely partially due to malignant T cells. Testicular DLBCL demonstrated high cytolytic score, in contrast to central nervous system DLBCL, indicating differences related to tumor site. Together, these data show that cytolytic score captures variation in cytolytic infiltrates across hematological malignancies and indicate that even in disease entities with generally low cytolytic activity, a subset of cases with abundant cytotoxic lymphocyte infiltration can be identified.

### IFNγ signature linked to cytolytic infiltration distinguishes lymphoma microenvironment from leukemias

To characterize cancers with abundant cytolytic infiltrate in more detail, we explored genes whose expression correlated with cytolytic score in each cancer type. As expected, genes positively correlated with cytolytic score were enriched in signatures reflecting T cell activation and inflammatory response, also confirmed at the protein level (Figures S2A and S2B and Table S2). To dissect genes expressed in cell types other than CTLs and NK cells, we contrasted the transcripts correlated with cytolytic score with the difference in expression between purified CTLs/NKs and the unsorted tumor samples (Figures 2A and S2C). We also investigated which normal cell types expressed the identified genes to define cell types co-infiltrating with cytolytic cells (Table S2). This analysis revealed strong correlations of genes expressed in monocytes and macrophages to cytolytic score both in B cell lymphomas and chronic leukemias (e.g. *CD14, R* > 0.6, FDR = 0.0), suggesting frequent coinfiltration of myeloid cells with cytotoxic lymphocytes (Figure 2B). In contrast, correlations of microenvironmental genes with cytolytic score were much more modest in AML, pre-B-ALL, T-ALL, and myelodysplastic syndrome (MDS) (e.g. *CD14 R* ≤ 0.3).

**Figure 2.**
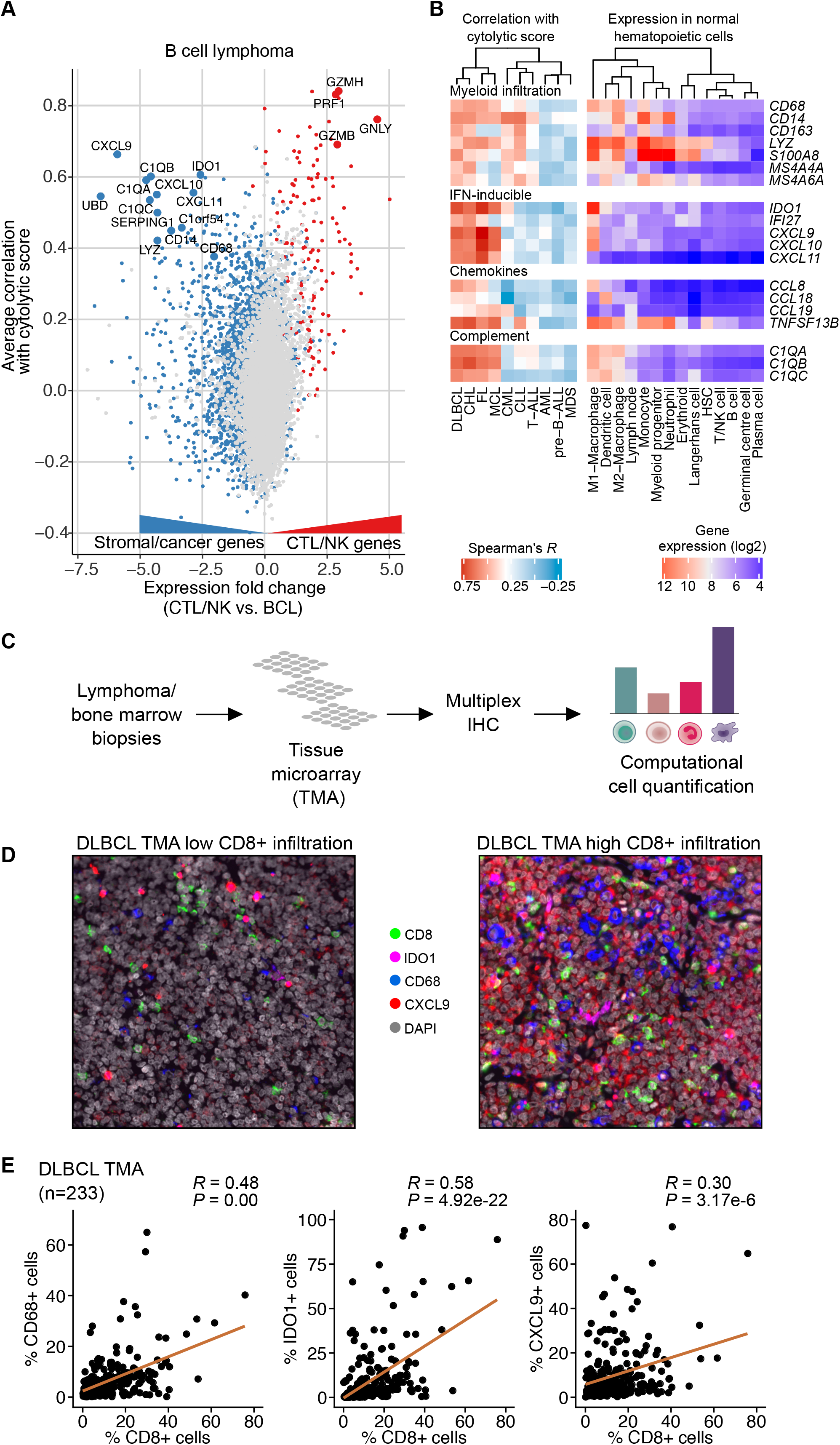
Distinct immune microenvironments associate with cytolytic infiltration in lymphoma and leukemia. **A**. Correlation of gene expression with CTL/NK abundance in B cell lymphoma. The average correlation between the expression of each gene and the cytolytic score in BCL samples (y axis) is compared to the fold change of the respective expression level between purified CTLs/NKs and bulk BCL transcriptomes (x axis). Genes more specific to CTLs/NKs are colored in red and genes more specific to other cell types in the tumor sample (stromal/cancer cells) are colored in blue. **B**. Heatmaps of correlation coefficients (Spearman correlation to cytolytic score) of selected genes across Hemap leukemia and lymphoma subtypes (left) and mean gene expression in normal cell types (right). **C**. Workflow of multiplex immunohistochemistry-based validation of protein-level expression of genes correlating with cytolytic score in lymphoma and leukemia microenvironments. **D**. Multiplex immunohistochemistry images of in representative DLBCL samples with low (left) or high (right) percentage of CD8+ cells (CTLs). CD8 (CTL marker), CD68 (macrophage marker), CXCL9, IDO1, and DAPI stainings are shown. **E**. Scatter plots comparing the percentage of CD8+ cells (CTLs) out of total cells with the percentages of CD68+ cells (macrophages), IDO1+ cells, and CXCL9+ cells out of total cells in the DLBCL IHC cohort (n=233). Spearman correlation coefficients and *P* values adjusted using the BH method are shown. See also Figure S2 and Table S2.

Characteristic microenvironmental genes associated with cytolytic infiltration in lymphomas included those encoding the immunosuppressive tryptophan-catabolizing enzyme *IDO1* (Munn and Mellor, 2016), T cell-recruiting chemokines highly expressed in proinflammatory M1-type macrophages *(CXCL9, CXCL10, CXCL11)*, and complement components expressed in macrophages and dendritic cells (DCs) *(C1QA, C1QB, C1QC)* (Figure 2B). The expression of *IDO1* and the CXCR3-ligand chemokines is known to be strongly induced by IFNγ (Groom and Luster, 2011; Spranger et al., 2013), suggesting evidence of a microenvironmental response to IFNγ associated with cytolytic infiltration. Expression of these genes in normal lymph nodes and leukemias was low (Figure 2B), indicating cancer-associated modulation of the lymphoma immune microenvironment.

To validate the distinct lymphoma microenvironment characterized by myeloid infiltration and IFN-y-induced expression signature, we analyzed tissue microarrays (TMAs) constructed from DLBCL and AML BM biopsies using multiplex immunohistochemistry (Figure 2C). Consistent with the gene expression data, CTLs (CD8+) correlated with macrophages (CD68+) and IDO1+ and CXCL9+ cells in DLBCL (Figures 2D and 2E). In AML, however, IDO1+ or CXCL9+ cells were generally sparse and did not correlate with CTLs, in concordance with the gene expression data (Figure S2D). CD68 did not correlate either with CTLs, but was likely expressed on a subset of blasts, potentially confounding correlations. Collectively, these data indicate general macrophage/monocyte infiltration associated with cytolytic cells and a distinct immunological tumor microenvironment in lymphomas characterized by IFNγ-responsive genes.

### Cytolytic infiltration is associated with driver alterations and molecular subtypes in AML and DLBCL

We next asked whether specific genetic alterations or molecular subtypes could be associated with increased abundance of cytotoxic lymphocytes. We first explored correlations of cytolytic score to somatic mutations and CNVs in the TCGA AML dataset (Table S3). Cytolytic score positively correlated with mutations in the *TP53* tumor suppressor gene (FDR < 0.00018), as well as deletions located in the long arm of chromosome 5 (Figure 3A). These alterations often co-occurred with complex cytogenetics and elevated genome fragmentation, whereas no correlation to mutation load was detected. In contrast, the common AML driver mutations *FLT3* and *NPM1* preferentially occurred in samples with low cytolytic activity. The samples with high cytolytic score were enriched in a cluster characterized by an MDS-like transcriptomic phenotype (FDR = 10^-6^, Fisher’s exact test) that we have previously identified (Pölönen et al., 2019) and mutations frequently found in MDS such as *RUNX1* and *ASXL1*, suggesting that a distinct MDS-like transcriptional signature may be associated with elevated cytotoxic lymphocytes in AML (Figures 3B-E).

**Figure 3.**
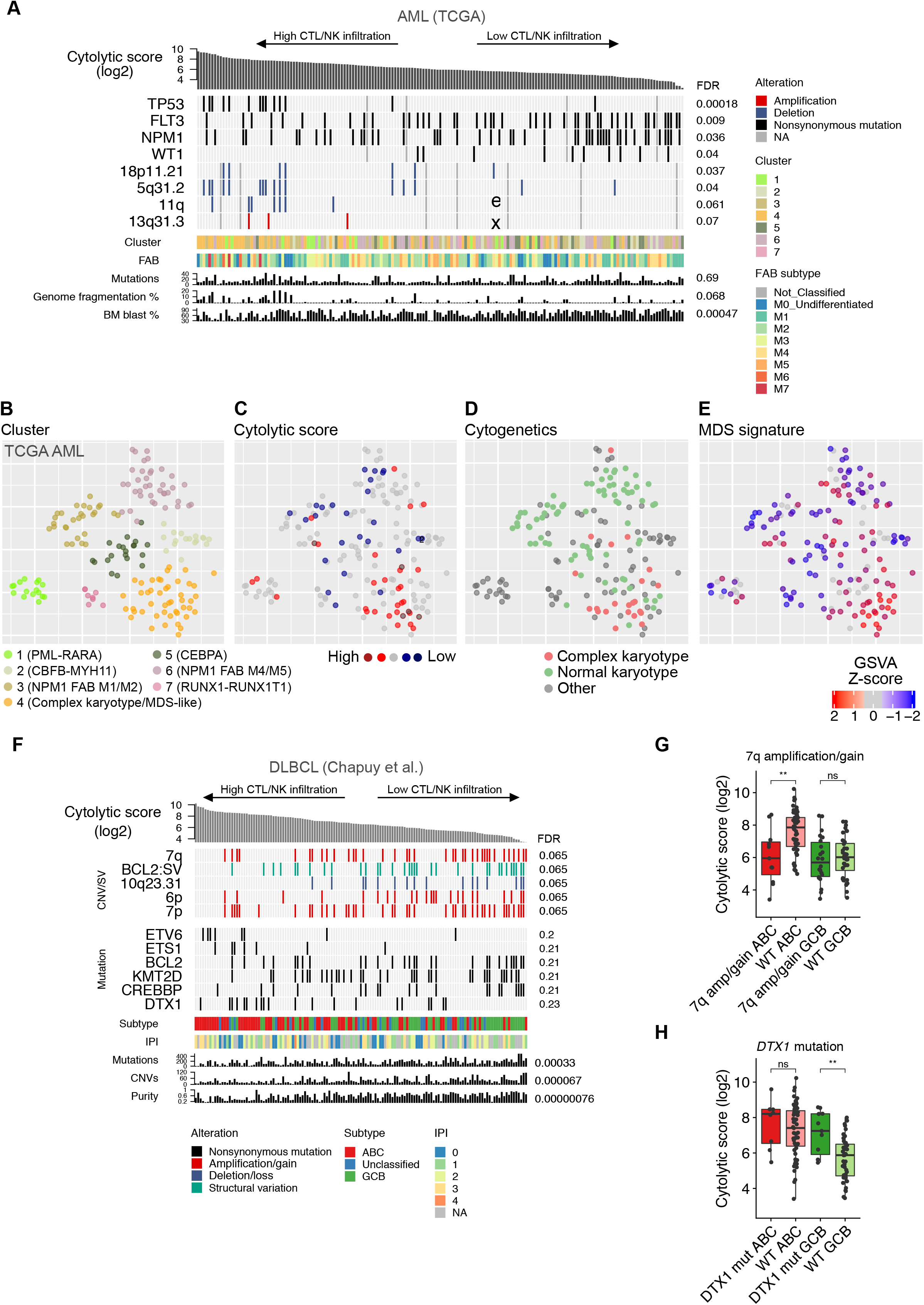
Genetic alterations associated with cytolytic infiltration. **A**. Top genetic alterations associated with cytolytic score in TCGA AML patients shown as an oncoprint, where columns corresponding to a sample are ranked by cytolytic score and different data values are plotted on rows. Discrete state/class is indicated as color (genetic alterations and sample categories) and continuous values are represented as barcharts (score or percentage values). FDR for correlations between cytolytic score and genetic alterations are shown. **B-D**. Visualization of TCGA AML samples using a t-SNE representation. Clusters, cytolytic score, cytogenetics, and MDS signature are colored on the t-SNE maps, respectively. Key characteristics of the clusters are annotated. **E**. Top genetic alterations associated with cytolytic score in DLBCL (data from GSE98588) patients are shown as in A. **F**. Comparison of cytolytic score in DLBCL between cases with 7q amplification and without (WT) stratified by molecular subtype. Nominal *P* values obtained using two-sided Wilcoxon rank sum test are shown. **G**. Comparison of cytolytic score in DLBCL between cases with *DTX1* mutations and without (WT) stratified by molecular subtype. Nominal *P* values obtained using two-sided Wilcoxon rank sum test are shown. See also Figure S3 and Table S3.

To validate the association of cytolytic infiltration to a transcriptomic subtype linked to complex cytogenetics and MDS-associated alterations, we identified matching transcriptional clusters in the Hemap AML and BeatAML datasets using a cluster-specific gene set enrichment approach (Mehtonen et al., 2019) (Figures S3A-D). The cases with high cytolytic score in Hemap AML and BeatAML were enriched in a cluster corresponding to the TCGA MDS-like cluster (FDR = 0.0044, Fisher’s exact test in BeatAML) with frequent complex cytogenetics and prior MDS cases. Cytolytic score correlated with diagnosis of AML with myelodysplasia-related changes (FDR = 0.05), further suggesting a link between an MDS-like/secondary AML subtype and increased cytolytic infiltration.

We next examined cytolytic score in relation to mutations and CNVs in DLBCL (Chapuy et al., 2018) (Table S3). *BCL2* translocations, which almost exclusively occur in GCB DLBCL, correlated negatively with cytolytic infiltration (FDR = 0.065, Figure 3F), consistent with the lower cytolytic score observed in this molecular subtype in Hemap (Figure 1H) and fewer CD8+ cells assessed by mIHC (Figure S3E). Several CNVs and GCB-associated mutations *(BCL2, KMT2D, CREBBP)* were negatively associated with cytolytic infiltration, whereas mutations in *ETV6, ETS1*, and *DTX1* showed positive correlations (FDR < 0.25). Cytolytic score correlated negatively with tumor purity assessed by ABSOLUTE (Carter et al., 2012), consistent with increased immune infiltrate linked to lower tumor cell fractions. Given the strong impact of the molecular subtype on cytolytic infiltrate, we analyzed both ABC and GCB subtypes separately to identify more direct associations to specific genetic alterations (Figures S3F and S3G). 7q amplifications were preferentially found in ABC with low cytolytic infiltration and GCB (Figure 3G). The correlation of *DTX1* mutations with cytolytic infiltration was even stronger in the GCB subtype alone compared to all DLBCL (FDR = 0.06, Figure 3H). Together, our data suggest that specific cancer cell-intrinsic genetic alterations and molecular subtypes are linked to cytotoxic infiltrate in both AML and DLBCL.

### Epigenetic modification of the HLA class II transactivator *CIITA* regulating antigen presentation in AML

Given the importance of effective antigen presentation for adaptive anti-tumor immune responses, we analyzed the expression of HLA genes to detect potential transcriptional downregulation. As the normal cellular counterpart of several hematological malignancies is an APC, the cancer cells could elicit T cell responses by presenting antigen in HLA class II molecules in addition to HLA I expressed in all nucleated cells. We constructed an HLA I score, comprised of *B2M, HLA-A, HLA-B*, and *HLA-C*, and an HLA II score, containing HLA II genes significantly upregulated in APCs (macrophages, DCs, B cells) compared to non-APCs and whose expression highly correlated with each other (Figures S4A and S4B and Table S4).

We observed a lower HLA I score in cells of the erythroid lineage, hematopoietic progenitors, and T cell acute lymphoblastic leukemia (T-ALL) compared to other cell populations (Figure 4A and Table S1). While the differences in HLA I expression were rather modest, HLA II expression varied more substantially. B cell malignancies, including pre-B-ALL, CLL, and B cell lymphomas had high HLA II score as expected by their APC origin, whereas in multiple myeloma (MM) HLA II was downregulated consistent with HLA II loss upon plasmacytic differentiation (Silacci et al., 1994) (Figures 4B and 4C and Table S1). In AML, specific transcriptomic clusters showed downregulated HLA II expression, including the acute promyelocytic leukemia cluster harboring PML-RARA fusion known to be characterized by low surface HLA-DR (Wetzler et al., 2003), and a cluster characterized by *NPM1* mutations and M1 or M2 FAB subtype. HLA II score correlated with HLA-DR surface expression in AML blasts measured using flow cytometry and paired RNA-seq, indicating that the HLA II score accurately reflects variation in surface HLA II protein levels (Figures 4D and S4C).

**Figure 4.**
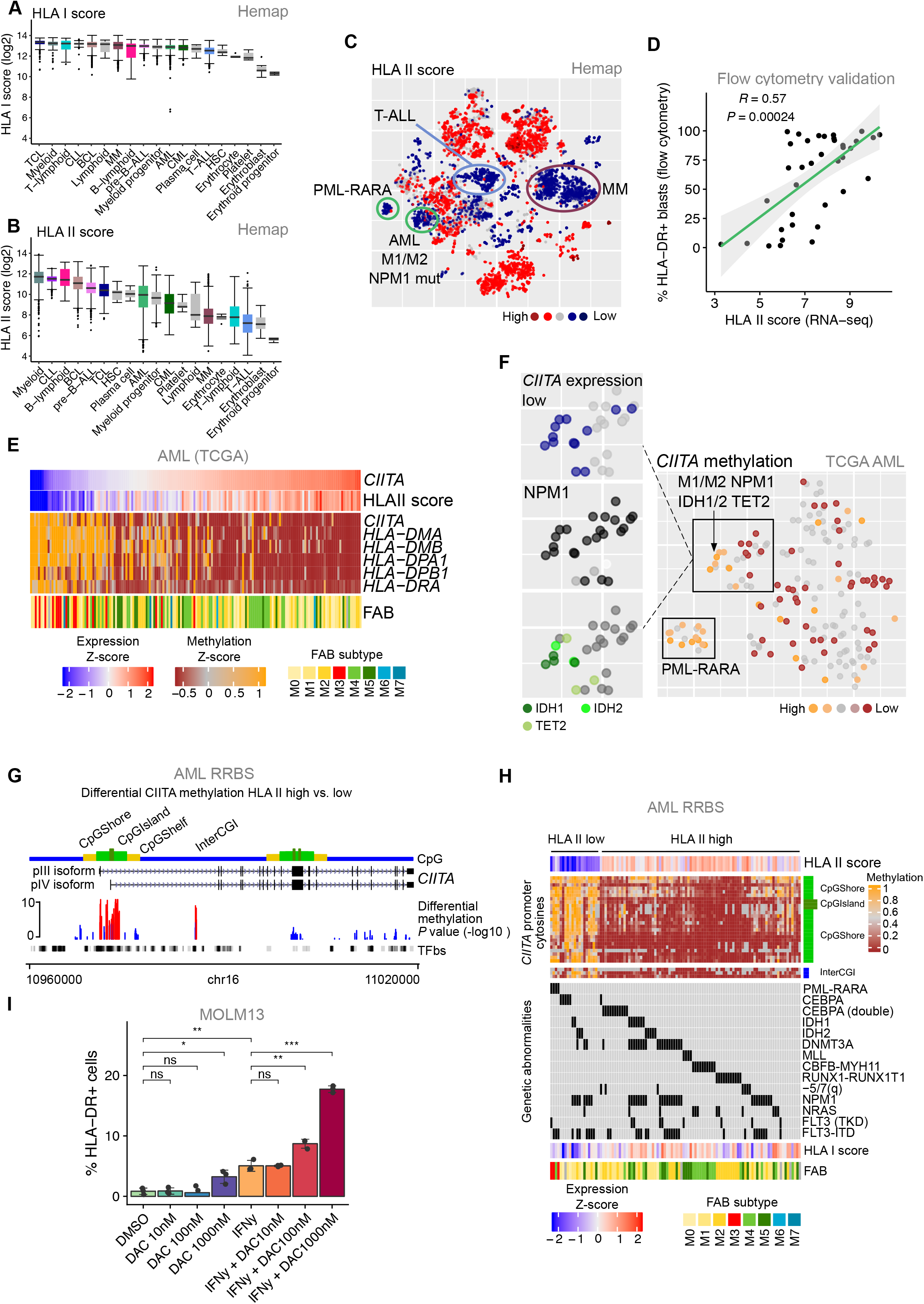
Expression of antigen-presenting HLA genes is linked to molecular subtypes and epigenetic regulation. **A**. HLA I score (log2 geometric mean of *B2M, HLA-A, HLA-B*, and *HLA-C*) is shown as boxplots comparing main cancer types and normal cell populations in Hemap. **B**. HLA II score (log2 geometric mean of *HLA-DRA, HLA-DRB1, HLA-DPA1*, *HLA-DPB1, HLA-DMA*, and *HLA-DMB)* is shown as in A. **C**. HLA II score colored on Hemap t-SNE map. Specific clusters with low HLA II score are highlighted with circles. **D**. Comparison of HLA II score and HLA II surface protein expression level in blasts in a validation cohort of AML BM samples (n = 37) profiled using both RNA-seq and flow cytometry for HLA-DR. **E**. *CIITA* expression, HLA II score, and methylation of *CIITA* and *HLA II* genes in TCGA AML data shown as a heatmap. **F**. Methylation of *CIITA* colored on the TCGA AML t-SNE map. *NPM1* and *IDH1, IDH2*, and *TET2* mutation status is labeled for cluster with low HLA II score. **G**. Differentially methylated cytosines (DMCs) in the *CIITA* region between samples with low and high HLA II score in GSE86952 ERRBS dataset. Histogram indicates the negative log10 *P* value of differential methylation at each cytosine, with red and blue colors indicating hypermethylated and hypomethylated cytosines in HLA II low samples, respectively. CpG areas, including CpG islands, CpG shores (< 2 kb flanking CpG islands), and CpG shelves (< 2 kb flanking outwards from CpG shores) are shown above *CIITA* exons belonging to isoforms pIII (lymphoid) and pIV (IFNγ-inducible). Transcription factor binding sites (TFbs) are shown below. **E**. Heatmap showing methylation of cytosines at *CIITA* regions significantly hypermethylated in the HLA II low group compared to high in the AML GSE86952 ERRBs dataset. 0 indicates no methylation and 1 indicates complete methylation. Patients (columns) are grouped by HLA II score and PML-RARA status. Rows correspond to cytosines at the CpG island, shores, and inter-CGI area (> 4 kb outwards from a CpG island) shown on the right. Major AML genetic alterations, HLA scores, and FAB classification are shown. **F**. Percentages of HLA-DR+ MOLM13 cells measured by flow cytometry after 72 h treatment with indicated concentrations of decitabine (DAC) and/or 10 ng/mL IFNγ. Dots indicate individual technical replicate wells. Data are shown for one of two independent experiments. *P* values are obtained using two-sided Wilcoxon rank sum test. See also Figure S4 and Table S4.

We next correlated molecular features, including mutations, CNVs, and DNA methylation in the TCGA AML cohort to the HLA II score to shed light on the molecular mechanisms leading to downregulation of the HLA II genes (Table S4). Expression of the HLA class II transactivator *CIITA* strongly correlated with the HLA II score (R = 0.84, FDR = 6.1 × 10^-43^, Figure 4E). However, methylation of promoter regions of *CIITA (R* = −0.54, FDR = 4.6 × 10^-10^, probe cg01351032) and several *HLA II* genes correlated negatively with the HLA II score (Figure S4D). *CIITA* methylation was enriched in transcriptomic clusters with low HLA II score corresponding to those identified in Hemap, harboring PML-RARA fusion or M1/M2 FAB subtype co-occurring with mutations in *NPM1* (Figures 4F and S4E). Upon closer examination, highest *CIITA* hypermethylation occurred in a cluster with mutations in the DNA methylation regulators *IDH1, IDH2*, and *TET2.* We observed a similar connection between epigenetic modifier mutations and low HLA II in the BeatAML dataset (Figure S4F), further suggesting a link between AML driver mutations and antigen presentation mediated by alterations in DNA methylation. In contrast, AML harboring CBFB-MYH11 or RUNX1-RUNX1T1 translocations were characterized by high HLA II and *CIITA* hypomethylation, and *RUNX1* mutations correlated with high HLA II score (FDR = 0.0013, TGCA and 2.9 × 10^-8^, BeatAML). In addition to AML, *CIITA* was methylated in T-ALL with low HLA II expression (Holling et al., 2004), suggesting potential epigenetic regulation of antigen presentation in different hematological cancer types (Figures S4G and S4H).

To validate the finding in an independent dataset, we analyzed differentially methylated cytosines (DMCs) in the *CIITA* promoter region between AML patients with high and low HLA II score using ERRBS data (Glass et al., 2017). This analysis demonstrated a differentially methylated region encompassing a CpG island and *CIITA* promoter III, active in lymphocytes, and the IFNγ-inducible promoter IV (Muhlethaler-Mottet, 1997) (Figures 4G and 4H). Further validating the observation, *CIITA* promoter methylation negatively correlated with HLA II score in AML cell lines (Figure S4G). In MOLM13 AML cells expressing low levels of HLA II with hypermethylation of the *CIITA* promoter, treatment with the hypomethylating agent decitabine partially restored HLA-DR surface expression and potentiated the HLA-DR induction by IFNγ, a known inducer of HLA II (Steimle et al., 1994) (Figures 4I and S4I). Taken together, these data show that AML cells may evade antigen presentation through transcriptional downregulation of *HLA II* genes, which is linked to *CIITA* methylation in distinct genetic and transcriptional subtypes of AML and across hematological cancers.

### Immune checkpoints are linked to cancer subtypes and genetic alterations

Immunomodulatory genes or immune checkpoints regulating T cell co-stimulation or NK cell activation represent important immunotherapy targets. We focused on ligands for T and NK cell co-stimulatory and co-inhibitory receptors and other immunomodulators to identify potentially targetable immune checkpoints in subtypes of hematological malignancies. We tested for enrichment of the immune checkpoints across cancer types and found distinct patterns of immunomodulatory genes in myeloid malignancies and mature B cell malignancies (Figure 5A and Table S5). The cancer samples also clustered together with their normal counterparts, suggesting that the cell-of-origin influences the repertoire of immunomodulatory genes expressed by cancer cells (Figure S5A).

**Figure 5.**
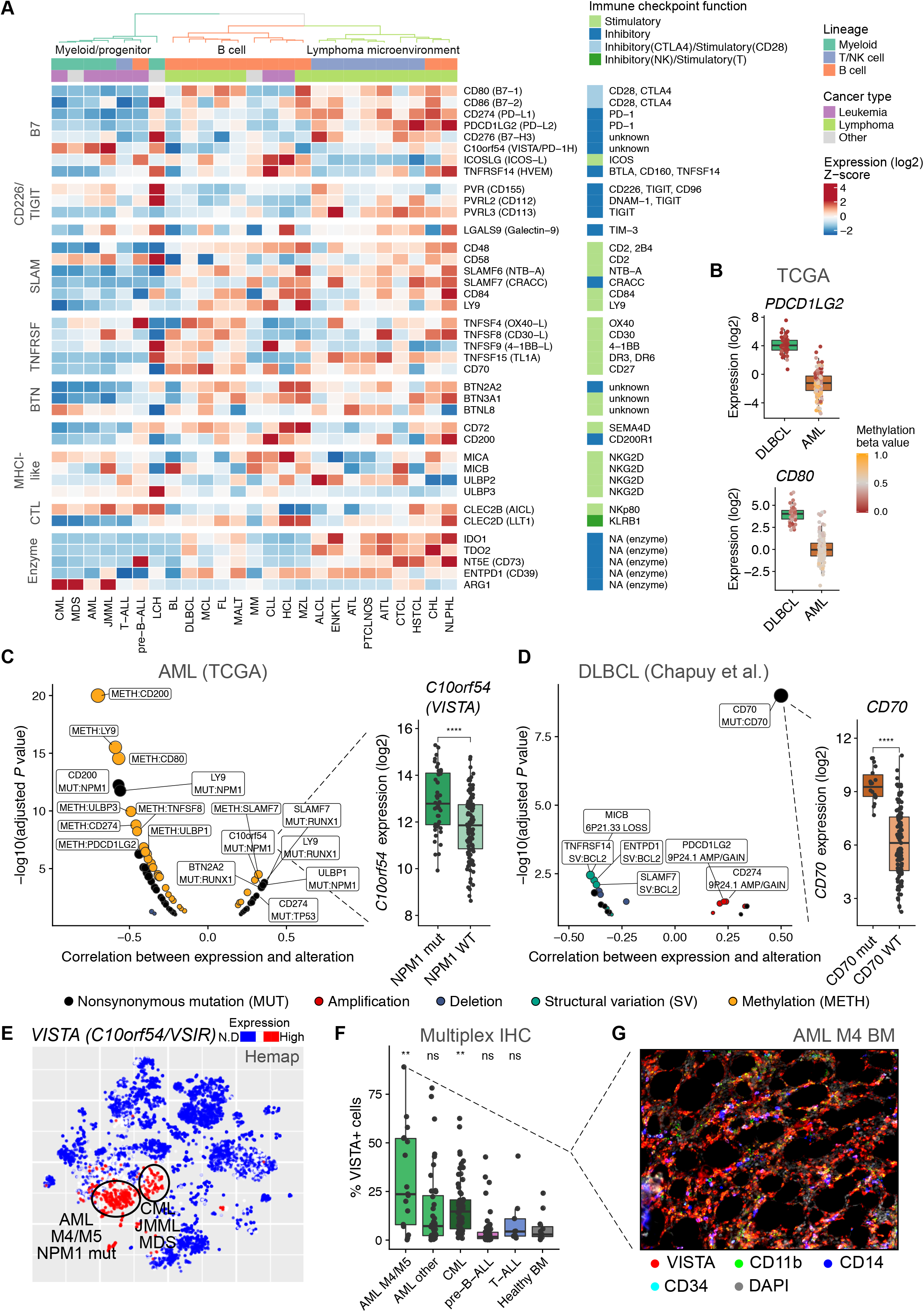
Immune checkpoints are linked to cancer subtypes and genetic alterations. **A**. Expression levels (Z-scores of median log2 expression) of co-stimulatory, co-inhibitory, and other immunomodulatory genes across Hemap cancer types shown as a heatmap. Sorted and unsorted samples are shown as separate categories. Rows and columns are clustered using Spearman correlation distance and Ward’s linkage. Corresponding receptors and the nature of the interaction is shown according to Table S5. **B**. Comparison of *PDCD1LG2* and *CD80* expression between TCGA DLBCL and TCGA AML. Dots indicating individual patients are colored by average methylation within 1 kb of the transcription start site. **C**. Volcano plot of correlations (Spearman) of immune checkpoint gene expression with genetic alterations and DNA methylation in TCGA AML. Dot size is proportional to the adjusted *P* value. *ULBP1* expression is compared between NMP1-mutated (mut) and wild type (WT) samples and the nominal *P* value obtained using two-sided Wilcoxon rank sum test is shown. **D**. Volcano plot of correlations (Spearman) of immunomodulatory gene expression with genetic alterations in DLBCL (GSE98588). Dot size is proportional to the adjusted *P* value. *CD70* expression is compared between CD70-mutated (mut) and wild type (WT) samples and the nominal *P* value obtained using two-sided Wilcoxon rank sum test is shown. **E**. Expression of *VISTA* colored on Hemap t-SNE map. Circles indicate clusters highly expressing *VISTA.* F. Percentages of VISTA-positive cells out of all BM cells in AML (n=57), CML (n=62), pre-B-ALL (n=51), T-ALL (n=9), and healthy BM (n = 11) tissue microarrays analyzed by quantitative multiplex immunohistochemistry. The *P* values indicate comparisons of leukemia types to healthy BM using two-sided Wilcoxon rank sum test. **G**. Multiplex immunohistochemistry of AML BM from a patient with M4 FAB subtype AML with high VISTA expression. VISTA, CD11b (myeloid marker), CD14 (monocyte marker), CD34 (blast/progenitor marker), and DAPI stainings are shown. See also Figure S5 and Table S5.

Myeloid malignancies, including AML, CML, JMML, and MDS, highly expressed *VISTA (C10orf54/VSIR/PD-1H)*, encoding an inhibitory T cell checkpoint of the B7 family (FDR < 10^-132^ AML compared to other cancers, Wilcoxon rank sum test). In addition to *VISTA, ARG1* encoding the immunosuppressive enzyme arginase represented another potential myeloid-specific immune evasion mechanism (FDR < 10^-84^ in AML). In contrast, the NK cell inhibitory receptor KLRB1 ligand *CLEC2D (LLT1)* (Figure S5B) and the T cell inhibitory butyrophilin *BTN2A2* were enriched in mature B cell malignancies. In addition to myeloid and B cell malignancies, lymphoma samples clustered together, characterized by elevated *CD274 (PD-L1), PDCD1LG2 (PD-L2), IDO* family enzymes, and *TNFSF15 (TL1A).* These genes were lowly expressed in the purified CD19+ lymphoma cells, but strongly in macrophages and DCs, suggesting microenvironmental origin of these genes (Figure S5A). Other cancer type-specific immune checkpoints included *PVRL3*, encoding a ligand for the inhibitory receptor TIGIT, in myeloma (FDR < 10^-156^) and T cell malignancies (Figure S5B), and *NT5E*, encoding the immunosuppressive adenosine-producing enzyme CD73 (Beavis et al., 2012), in pre-BALL (FDR < 10^-303^). Together, these results suggest cancer type-specific immune checkpoints in hematological malignancies, such as *VISTA* in myeloid malignancies.

To investigate potential mechanisms of immunomodulatory gene regulation leading to the observed expression patterns, we examined correlations with DNA methylation in AML and DLBCL in the TCGA dataset (Table S5). Comparison of AML and DLBCL revealed differential methylation at gene promoters linked to the cancer type-specific expression of several immunomodulators, such as *PDCD1LG2* (FDR = 4.4 × 10^-28^ for differential methylation and 4.3 × 10^-55^ for expression between AML and DLBCL) and *CD80* (FDR = 1.3 × 10^-62^ and 1.0 × 10^-47^) (Figures 5B, S5C, and S5D). In addition to variation between cancers, promoter methylation also correlated with expression of immunomodulators within cancer types. In AML, promoter methylation correlated negatively with expression of *Cd200, CD274*, the NKG2D ligands *ULBP1* and *ULBP3*, and *PDCD1LG2* and *CD80* (*R* ≤ – 0.4, FDR < 10^-6^, Figure 5C). Similarly in DLBCL, expression of *ULBP1* and *CD200*, among other genes, correlated negatively with promoter methylation, suggesting that DNA methylation contributes to variation in immune checkpoints across cancer types (Figure S5E).

In addition to epigenetic modification, several genetic driver alterations were linked to distinct immune checkpoints, suggesting potential immune evasion strategies. In AML, *NPM1* mutations were linked to elevated expression of *VISTA* (FDR = 0.00041) and the NKG2D ligand *ULBP1* (FDR = 0.00027, Figures 5C and S5F). RUNX1-mutated AML highly expressed the B cell-associated *BTN2A2, SLAMF7*, and *LY9* in addition to HLA II, suggesting that the lineage infidelity and B lineage transcriptional program induced by *RUNX1* mutations influences also co-inhibitory signaling by AML cells (Silva et al., 2009). *TP53* mutations were linked to higher *CD274 (PD-L1)* expression, potentially related to increased cytolytic activity. In DLBCL, the co-stimulatory *CD70* was often mutated when highly expressed (FDR = 10^-9^, Figure 5D), suggesting evasion from the T cell stimulatory interaction through somatic mutations. Other alterations potentially enabling immune evasion included downregulation of *MICB*, encoding an activating ligand for the NKG2D receptor, through 6p21.33 losses containing *MICB*, and 9p24.1 amplifications or gains associated with elevated expression of PD-1 ligands (Figure S5G). Thus, genetic alterations contribute to variation in immunomodulatory gene expression in hematological malignancies such as AML and DLBCL.

Finally, to validate the cancer type-specific immune checkpoints at the protein level, we performed mIHC on BM biopsies focusing on VISTA which we identified enriched in myeloid malignancies. In Hemap, *VISTA (C10orf54)* expression was strongly enriched in a cluster representing NPM1-mutated AML with M4/M5 FAB subtype, and a cluster comprising MLL-rearranged cases, suggesting association of *VISTA* expression with monocytic differentiation of leukemic cells (Figure 5E). Quantitative mIHC confirmed elevated VISTA in monocyte-like AML and CML BM compared to lymphoid malignancies or healthy controls (Figures 5F, 5G, and S5H). Together, our data suggest that several immune checkpoints such as VISTA are expressed in a cancer type-specific fashion and may be influenced by DNA methylation or driver alterations.

### Frequent expression of cancer-germline antigens in multiple myeloma

To evaluate potential targets of the adaptive cytotoxic immune response, we investigated cancer-germline antigens (CGAs) that can be readily identified from transcriptomic data but have not been systematically studied in hematological malignancies. CGAs are expressed only in the immune-privileged germ cells among healthy tissues, but aberrantly activated in cancers. We integrated the Genotype-Tissue Expression (GTEx) project (Mele et al., 2015) and Hemap data to define genes with a cancer-germline expression pattern in hematological malignancies by first selecting genes expressed in testis but not in other human tissues using GTEx and then requiring the genes to be expressed in 5% hematological cancers but not in normal hematopoietic cells (Figure 6A). Using these stringent criteria, we recovered 27 CGA genes. Most of the genes are included in the CTdatabase (Almeida et al., 2009), which however contains several genes whose expression is not testis-restricted (Figure S6A).

**Figure 6.**
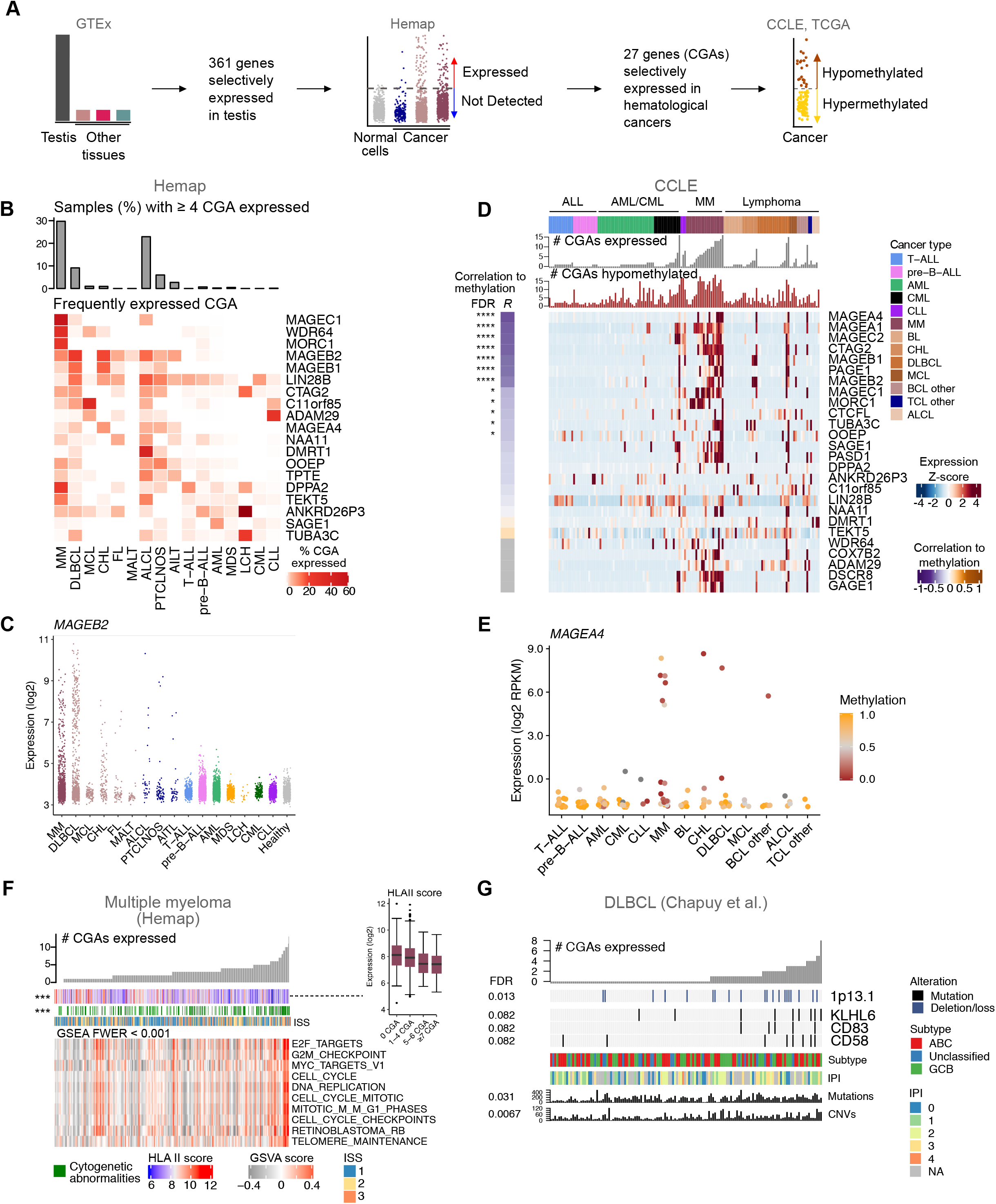
Cancer-germline antigen expression is frequent in multiple myeloma and linked to DNA methylation. **A**. Schematic of identification of genes with cancer-germline expression pattern. **B**. Expression of CGAs across hematological malignancies (Hemap). Color indicates percentage of samples from each cancer types expressing a given antigen. **C**. Expression of *MAGEB2* across Hemap cancer types and healthy samples. Examples of expression of other CGAs are shown in Figure S6C. **D**. Expression of CGAs across cell lines of hematological malignancies (CCLE) shown as a heatmap. Numbers of expressed (> 0.5 RPKM) and hypomethylated (< 0.5 average methylation) CGAs in each cell line are indicated as bars above the heatmap. Correlation (Spearman) of gene expression with average methylation and the corresponding FDR is shown on the right. Cell lines within cancer types are ordered by number of expressed CGAs and genes are ordered by correlation coefficient between gene expression and average methylation. **E**. Expression of *MAGEA4* in CCLE hematological cancer subtypes. Dots indicating individual cell lines are colored by average methylation value, where 0 indicates no methylation and 1 indicates complete methylation. **F**. Gene sets correlated with the number of expressed CGAs in MM. Heatmap shows GSVA scores of gene sets for each patient. **G**. Genetic alterations correlated with the number of expressed CGAs in DLBCL (Chapuy *et al*.). See also Figure S6 and Table S6.

Strikingly, across hematological cancers, CGAs were most frequently expressed in multiple myeloma (MM) (Figures 6B and S6B). One third of MM patients expressed more than four CGAs. Both B and T cell lymphomas also showed frequent CGA expression, whereas CGAs were largely transcriptionally silent in leukemias. Several CGAs were expressed in a cancer type-specific manner, including *MAGEC1, MORC1, DPPA1, COX7B2, PAGE1*, and *GAGE1* in MM, *ADAM29* in CLL (Vasconcelos et al., 2005), *SAGE1* in AML, *DMRT1* in anaplastic large-cell lymphoma (ALCL), and *MAGEB2* and *MAGEB1* in DLBCL (Figures 6C and S6C).

To understand mechanisms leading to aberrant CGA expression in MM and lymphomas, we studied whether alterations in DNA methylation are associated with activated CGA transcription. CGAs were frequently hypomethylated and expressed in myeloma cell lines in CCLE, consistent with primary myeloma samples (Figures 6D and 6E). Similarly, expression of the most frequent CGAs in DLBCL in patients in the TCGA dataset, *MAGEB1* and *MAGEB2*, was linked to hypomethylation at probes located near the transcription start site (Figures S6D and S6E).

We next explored transcriptional and genetic signatures correlated with the number of expressed CGAs. In Hemap MM, CGA expression was associated with gene sets reflecting cell cycle and MYC targets, implying that highly proliferative cancers frequently express multiple CGAs (Figure 6F). Furthermore, the presence of cytogenetic abnormalities was linked to increased CGA expression, whereas HLA II score correlated negatively with the number of expressed CGAs. In DLBCL, number of CGAs correlated with mutation and CNV load (FDR < 0.05) as well as specific alterations such as 1p13.1 deletions containing *CD58* (FDR = 0.02) and *CD58* mutations (FDR = 0.1), providing a potential immune evasion mechanism for CGA-expressing cancers through disruption of the CD2-CD58 interaction with T cells (Figure 6G). In the ABC subtype, a mutational signature including 6q deletions and *MYD88, HLA-A*, and *ETV6* mutations (FDR < 0.15), resembling cluster 5 (Chapuy et al., 2018), was enriched in cases expressing multiple CGAs (Figure S6F), suggesting a link between a distinct genetic subtype and activation of germline-restricted genes. Cytolytic score and gene sets reflecting inflammatory response were interestingly downregulated in ABC DLBCL expressing multiple CGAs (Figure S6G). In the GCB subtype, 1p13.1 deletions and *KLHL6* mutations correlated with CGA expression, similarly as in all DLBCLs (FDR < 0.08). Together, these data suggest that CGA expression is activated in myelomas and lymphomas harboring genomic aberrations or distinct genetic alterations associated with immune evasion, often involving promoter hypomethylation.

### Immunological features are associated with survival

Finally, we aimed to delineate how immunological features are associated with overall survival. To comprehensively profile the prognostic associations of immune properties, we obtained survival models using elastic net Cox proportional hazards modeling in DLBCL, MM, and AML where multiple datasets with outcome data were available in Hemap (Table S7). Tested features included cytolytic score, HLA scores, number of expressed CGAs and individual CGAs, immunomodulatory genes, microenvironmental genes linked to cytolytic infiltrate, as well as established clinical risk scores, International Prognostic Index (IPI) in DLBCL and International Staging System (ISS) in MM. We used Hemap datasets for training the models (Figures S7A and S7B) and validated the results in independent test cohorts. The established models remained prognostic for overall survival in independent external validation datasets (Figures 7A and 7B), indicating that the identified associations of immune features to survival are robust. Although clinical risk scores were strong predictors of survival both in DLBCL and MM as expected, immunological features significantly improved outcome predictions in both cancer types, further stratifying patients within existing risk groups, including the cell-of-origin subtype and IPI in DLBCL and ISS in MM (Figures S7C and S7D).

**Figure 7.**
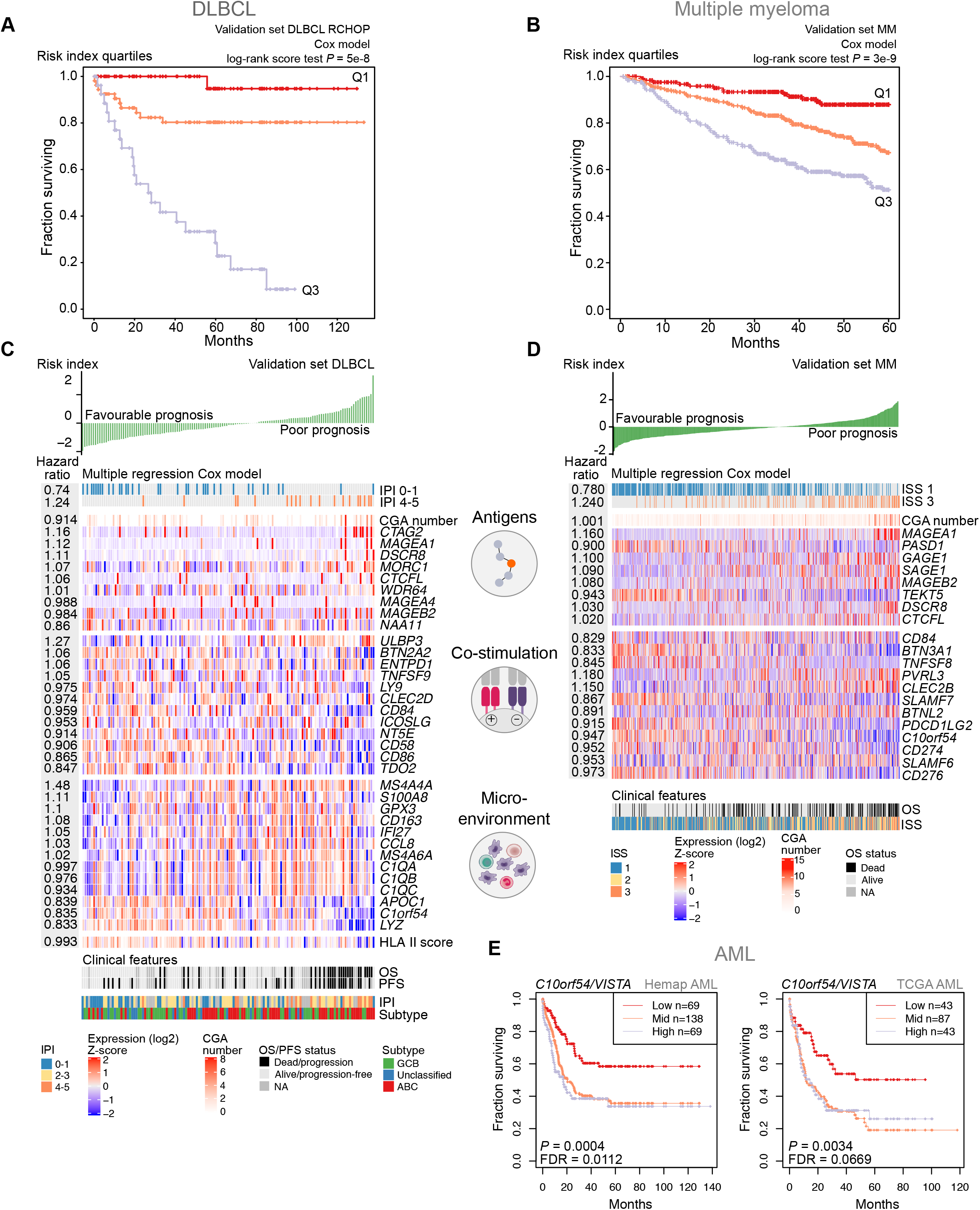
Immunological properties are associated with survival. **A**. Kaplan-Meier curves of overall survival for DLBCL patients stratified by the risk index of the immunological risk model (shown in Figure 7C) in the validation set (GSE98588). Patients were divided into three groups based on quartiles and *P* value was obtained using the score (log-rank) test. **B**. Kaplan-Meier curves of overall survival, as in A, for MM patients stratified by the risk index of the immunological risk model (shown in FIgure 7D) in the validation set (GSE16716 and GSE24080). **C**. Heatmap depicting features included in the DLBCL immunological risk model in the validation set (GSE98588). Patients are ordered by the risk index and hazard ratio (HR) for each feature is shown on the left. Features are grouped based on their function and clinical features are shown below the model heatmap. **D**. Heatmap depicting features included in the MM immunological risk model in the validation set (GSE16716 and GSE24080). Patients are ordered by the risk index and hazard ratio (HR) for each feature is shown on the left. Features are grouped based on their function and clinical features are shown below the model heatmap. **E**. Kaplan-Meier curves of overall survival for AML patients stratified by *C10orf54/VISTA* expression in Hemap and TCGA AML. Patients were divided into three groups based on quartiles and *P* value was obtained using the Wald test. See also Figure S7 and Table S7.

In DLBCL, certain monocyte/macrophage genes correlated with cytolytic score, such as *LYZ* and *APOC1*, strongly associated with better overall survival according to the risk model and also in univariate analysis (Figures 7C, S7E, and S7F). However, several macrophage-associated genes, including *C1QA, C1QB, C1QC, CD163*, and *MS4A6A*, were linked to worse survival, suggesting that distinct types or states of infiltrating myeloid cells characterized by these genes may have opposing impact on outcomes. The co-stimulatory genes *CD58* (CD2 ligand) and *CD86* (CD28/CTLA-4 ligand) were associated with improved outcomes. HLA II expression was also associated with survival benefit, consistent with previous findings (Lenz et al., 2008).

In MM, the butyrophilin *BTN3A1* expressed in B cell malignancies was linked to better overall survival, reflected also in univariate analysis (Figures 7D, S7G, and S7H). Several immune checkpoint receptor ligands associated with superior survival, including *CD274 (PD-L1), PDCD1LG2 (PD-L2), VISTA (C10orf54)*, and *CD276.* However, the TIGIT ligand *PVRL3 (CD113)*, highly expressed in MM compared to other cancers, was linked to poor survival. The expression of several CGAs was linked to worse outcome in both MM and DLBCL, suggesting increased CGA expression in more aggressive or advanced cancers.

In AML, we found establishing a risk model generalizable across datasets challenging, likely due to heterogeneity in the composition of the studied patient cohorts. However, we were able to identify individual genes with robust survival associations across multiple datasets (Table S7). Elevated *VISTA (C10orf54)* expression was associated with adverse outcomes in both Hemap and TCGA AML datasets (Figure 7E). Together, these findings show that immunological properties of hematological cancers have complex associations with survival.

## DISCUSSION

Understanding the determinants that shape the immunological landscape in cancer subtypes could enable more precise development of immune intervention approaches. We explored large-scale genomic datasets to uncover factors explaining immunological heterogeneity in hematological malignancies. We validated the discoveries in independent datasets and using orthogonal methods including multiplex immunohistochemistry and flow cytometry, lending robustness to the findings. The data suggest that both microenvironmental properties and cancer cell-intrinsic genetic and epigenetic features are associated with cytotoxic immune response, expression and presentation of antigens, and immune checkpoints. Our findings thus underline the importance of integrating data of genetic and epigenetic aberrations as well as the tumor microenvironment for a complete understanding of factors that may impact immunotherapy responsiveness.

We found consistently higher cytolytic score in lymphomas compared to other hematological malignancies, suggesting a higher ratio of cytotoxic lymphocytes to cancer cells in the lymphoma microenvironment. Increased cytolytic activity within lymphomas was associated with monocyte/macrophage-derived and IFNγ-inducible genes. A similar IFNγ-related profile has been shown to predict clinical response to PD-1 blockade (Ayers et al., 2017). Specific types of macrophages, monocytes, and DCs expressing these genes were recently identified using single-cell transcriptomics in melanoma (Li et al., 2018), corroborating our cell type inference and suggesting that similar cell types may infiltrate solid tumors and lymphomas. A distinct microenvironmental response might thus influence efficacy of immunotherapies in lymphomas as opposed to acute leukemias, where we were unable to detect similar microenvironmental signatures associated with cytolytic activity.

We identified cancer cell-intrinsic genetic alterations linked to cytotoxic infiltration, including *TP53* mutations, 5q deletions, and complex karyotype in AML. These genetic aberrations have been demonstrated to co-occur in elderly AML patients with dismal prognosis (Rücker et al., 2012). Both negative (Rooney et al., 2015) and positive (Thorsson et al., 2018) associations between *TP53* alterations and immune infiltration have been observed in other cancers. In DLBCL, *DTX1* mutations marked an immune-infiltrated group of especially GCB lymphomas. Cytolytic infiltration was linked to distinct molecular subtypes, including AML with myelodysplasia-related changes and ABC DLBCL, suggesting that the molecular phenotype of cancer cells may influence the immune infiltrate together with genetic alterations.

Downregulation of HLA class II genes was associated with hypermethylation of the transactivator *CIITA*, potentially resulting in defective antigen presentation to T helper lymphocytes. As HLA loss has been linked to AML relapse after allo-HSCT (Vago et al., 2009), low expression level already at diagnosis could restrict CD4+ T helper cell-mediated recognition both during an autologous immune response and in the allogeneic setting. Moreover, *CIITA* hypermethylation could be responsible for the transcriptional downregulation of HLA II upon relapse after allo-HSCT (Christopher et al., 2018). Given the reversible nature of epigenetic silencing as demonstrated by combined hypomethylating and IFNγ treatment of AML cells, reversal of promoter methylation could potentially augment HLA II-dependent immunity. Of interest, combining PD-1 blockade immunotherapy with hypomethylating agents has demonstrated efficacy in AML patients (Daver et al., 2018).

Several co-inhibitory immune checkpoints were expressed in a cancer type-specific fashion. Targeting different inhibitory interactions might thus be required for maximizing immunotherapy benefit in each disease. *VISTA* emerged as a novel checkpoint enriched in myeloid malignancies, including CML, MDS, JMML, and particularly monocytic, NPM1-mutated AML. *VISTA* expression was also linked to inferior outcomes in AML. *VISTA* is expressed in monocytes and neutrophils in healthy hematopoiesis (Flies et al., 2014), and could potentially be utilized by cancer cells of these lineages for immune evasion. VISTA has been implicated as a potential novel immunotherapy target in some solid tumors, such as prostate and pancreatic cancer and mesothelioma (Blando et al., 2019; Gao et al., 2017; Hmeljak et al., 2018). We also identified both genetic and epigenetic factors impacting immunomodulatory gene expression, such as high expression of *VISTA* in NPM1-mutated AML and copy number losses of *MICB*, encoding a ligand for the activating T/NK cell receptor NKG2D, thus providing additional potential layers of regulation to the cancer type-specific expression patterns.

CGA expression was more frequent in multiple myeloma and B and T cell lymphomas compared to other hematological malignancies. Although comparison to other cancers has not been previously performed, CGA expression and anti-CGA immune responses have been demonstrated in MM (Atanackovic et al., 2007; van Duin et al., 2011). Higher mutation loads have been described in MM and DLBCL compared to other hematological malignancies (Alexandrov et al., 2013), suggesting higher immunogenicity in these diseases conferred both by neoantigens and CGAs. CGA expression correlated with promoter hypomethylation and poor prognosis both in MM and DLBCL. Consistently, progression from monoclonal gammopathy with undetermined significance to advanced MM has been linked to global hypomethylation (Heuck et al., 2013). Thus, treatment of advanced myelomas could potentially benefit from immunotherapies leveraging the high number of expressed antigens, such as vaccines or T cell receptor therapies tailored for most common CGAs. CGA expression has been associated with resistance to CTLA-4 blockade (Shukla et al., 2018) and, in contrast, with response to PD-1 inhibition (Saghafinia et al., 2018), suggesting also relevance for patient stratification for immune checkpoint blockade therapies. In contrast, CGA expression was sparse due to hypermethylation in leukemias, where treatment with hypomethylating agents could be used increase antigenicity.

The immunological risk models revealed complex links between immunological features and patient survival, and highlighted potential immune properties that could be targeted to improve outcomes. In AML, the unfavorable prognosis linked to high *VISTA* expression further underlines VISTA as a potential target for novel immune checkpoint blockade approaches. Distinct subsets of monocyte/macrophage genes correlated with CTL/NK infiltration displayed diverging survival associations in DLBCL. Different cell populations marked by these genes co-infiltrating with cytotoxic lymphocytes may modulate the resulting immune response thus influencing outcomes, possibly explaining the lack of favorable prognostic survival association of cytolytic score itself. Although the cohorts studied here have received immunomodulatory treatments such as rituximab in DLBCL and thalidomide in MM, correlating immune signatures to outcomes of novel immunotherapies such as immune checkpoint blockade or CAR T cell therapy may reveal patterns distinct from those highlighted here.

Our approach of estimating immune cell composition from bulk gene expression data is limited in the analysis of rare cell types and normal immune cells transcriptionally resembling cancer cells, which is often the case in hematological malignancies. We anticipate that single-cell transcriptomics studies will further illuminate the association between infiltrating immune cell types and their transcriptional programs, such as those regulating antigen presentation or co-stimulatory signaling. Moreover, the genotypic-immunophenotypic associations identified from genomic data are unable to yield mechanistic insights into causal relationships between tumor genetics and immune states. We envision that the results presented here can guide further experimental investigation into the underlying tumor characteristics that modulate inter-tumor heterogeneity in immune landscapes.

In summary, our integrative analysis provides evidence of genomic and microenvironmental factors associated with variation in the immune contexture between different tumors. The findings of this study highlight the need to integrate genetic, epigenetic, and transcriptomic data of different aspects of the immune landscape to understand potential determinants of responsiveness to cancer immunotherapies.

## MATERIALS AND METHODS

### Patients

#### RNA sequencing and flow cytometry

Bone marrow (BM) aspirates from 37 AML patients were collected at diagnosis after signed informed consent from each patient (permit numbers 239/13/03/00/2010, 303/13/03/01/2011, Helsinki University Hospital (HUH) Ethics Committee) in accordance with the Declaration of Helsinki.

#### Tissue microarrays (TMA)

We collected diagnostic BM biopsies from AML (n=66), B-ALL (n=54), T-ALL (n=14), and CML (n=62) patients treated in the Department of Hematology, HUH between 2005-2015, and DLBCL biopsies treated at the Department of Oncology, HUH (n=233). In addition, BM biopsies taken in 2010 from subjects due to persistent abnormal leukocyte, erythrocyte, or platelet count and without diagnosis of hematological malignancy, chronic infection, nor autoimmune disorder in six years of follow-up were included as controls (n=11). Study subjects gave written informed research consent to the study and to the Finnish Hematology Registry. The study complied with the Declaration of Helsinki and the HUH ethics committee (permit number 303/13/03/01/2011). Fresh BM biopsies and lymphoma samples were formalin-fixed and paraffin-embedded (FFPE) in the Department of Pathology, HUSLAB and stored at the Helsinki Biobank at HUH. TMA blocks were constructed from up to four 1 mm cores from representative regions of tumor samples. RNA sequencing-defined molecular subtypes were available from a subset of DLBCL TMA patients (Reddy et al., 2017).

### Cell line

The MOLM13 cell line established from the peripheral blood of a 20-year-old man with AML was obtained from the Deutsche Sammlung von Mikroorganismen und Zellkulturen GmbH (DSMZ). Cells were cultured in RPMI-1640 (Lonza) with 10% FBS, 2 mM L-glutamine, 100 U/mL penicillin, and 100 μg/mL streptomycin (R10). The cell line was authenticated using GenePrint10 System (Promega) and confirmed to have an overall identity estimate of 100% at all 18 tested alleles.

### Processing of genome-wide multilevel data

Each dataset and sample numbers used in the analysis are listed in Figure S1.

#### Hemap

The Hemap dataset includes 9,544 gene expression profiles collected across several studies from the Gene Expression Omnibus (GEO) database described in (Pölönen et al., 2019). The data, curated sample annotations and disease categories are available at http://hemap.uta.fi. Briefly, these data represent microarray data from the commonly used hgu133Plus2 platform that were processed using the RMA probe summarization algorithm (Irizarry et al., 2003) with probe mapping to Entrez Gene IDs (from BrainArray version 18.0.0, ENTREZG) and a bias-correction method (Eklund and Szallasi, 2008) to generate gene expression signal levels. To ensure representative samples for the immunological analyses, we performed filtering on the original dataset. Treatments that induced cell differentiation or activation of normal cells were kept, while other *ex vivo* treatments (617 samples), non-malignant cells from patients (417), or clusters representing only a single study (>90% samples with the same study ID, 38 samples) were excluded from the analysis performed here, resulting in a dataset of 8,472 samples. We also distinguished sorted (using FACS, isolated by magnetic beads using either positive or negative selection, or microdissected), unsorted, and cell line samples based on sample descriptions. Finally, clinical information was added when available (survival from GSE10358, GSE10846, GSE11877, GSE12417, and GSE14468; progression-free survival, sex, age, race, and tumor cell contents from GSE13314, GSE10846, GSE24080, and GSE19784). All annotations for the samples used are provided in Table S1.

#### TCGA

Processed data were retrieved into feature matrices for AML and DLBCL, containing expression, mutation, CNV, methylation, and clinical data, using TCGA Feature Matrix Pipeline and fmx-construction.sh command, available at https://github.com/cancerregulome/gidget. For representing the data at individual gene loci, level 3 RSEM RNA-seq and methylation data for each TCGA AML and DLBCL sample was obtained from firehose GDAC, run stddata_2015_11_01. Methylation data were annotated using FDb.InfiniumMethylation.hg19 R package to assign probes at TSS.

#### CCLE

RNA-seq read counts (RPKM) dated 2018.05.02, RRBS methylation data for CpG islands and TSS 1 kb dated 2018.06.14, and cell line annotations dated 2012.10.18 were downloaded from: https://portals.broadinstitute.org/ccle/data.

#### Other multilevel datasets

Affymetrix Human Genome U133 Plus 2.0 gene expression datasets (DLBCL: GSE98588 and AML: GSE6891) were normalized using affy 1.52.0 (Gautier et al., 2004) RMA and gene expression values obtained using Brainarray v18 probe mapping. Mutations, chromosomal rearrangements, and clinical and sample characteristics were obtained from Supplementary Tables 2-5 for the DLBCL study (Chapuy et al., 2018). Clinical data, mutations, and sample annotations were obtained from Supplementary Tables 1-2 for AML (Glass et al., 2017).

Methylation data from pre-B-ALL and T-ALL samples in GSE49031 (processed beta values for each probe) were used in the analysis of *CIITA* expression and methylation *(CIITA* probe cg04945379).

BeatAML mutation, clinical, and sample annotation data were downloaded from source data (from Supplementary Table 3) (Tyner et al., 2018). The RNA-seq count matrix was obtained from the authors. Genes with log2 cpm level > 1 in over 1% of samples were voom transformed and quantile normalized using limma (Ritchie et al., 2015). Mutation status was assigned based on exome sequencing and clinical sequencing data. Only bone marrow samples were used in the statistical analysis of immunological features.

### Sample stratification based on gene expression profiles

Molecular subtypes were identified from the Hemap, TCGA AML, and BeatAML datasets using an data-driven approach as previously described (Mehtonen et al., 2019). Briefly, the Barnes-Hut approximated version of t-SNE implementation (BH-SNE) (16) was used with 15% most variable genes to perform dimensionality reduction. Kernel density-based clustering algorithm known as mean-shift clustering (Cheng, 1995) with bandwidth parameter set to 1.5 (subsets of data, one cancer type) or 2.5 (all data) was used (LPCM package in R) to cluster the data following the dimensionality reduction. To identify corresponding clusters in different datasets of the same cancer type, similarity in sample clustering between datasets was evaluated in a data-driven manner using GSVA (Hänzelmann et al., 2013) enrichment scores as previously described (Mehtonen et al., 2019). Briefly, top 20 positively and negatively correlated genes per cluster were used to identify similar clusters in a new dataset with significant enrichment for cluster specific genes.

### Statistical analysis using discrete gene expression features

For individual genes, discrete categories (high, low, and not detected) were assigned based on mixture model fit as described previously for Hemap and DLBCL GSE98588 datasets (Pölönen et al., 2019). Briefly, Gaussian finite mixture models were fitted by expectation-maximization algorithm (R package mclust version 4.3). The model was chosen by fitting both equal and variable variance models and ultimately choosing the model which achieved a higher Bayesian Information Criterion (BIC) to avoid overfitting. To assure minimal amount of misclassifications of data samples to discrete categories, three additional rules were implemented. First, if the uncertainty value from the model was above 0.1, value of 0 was assigned to denote low level. Secondly, log2 expression values lower than 4 or higher than 10 were assigned to a value −1 and 1, respectively. Thirdly, genes without clear background distribution (gene is always expressed), or if over 90% of the samples had uncertain expression based on the model classification, categories were re-evaluated. If >60% of the uncertain samples had expression above or below 6, categories were assigned as 1, and −1, respectively. For binary classification, values −1 and 0 were merged as low/not detected expression and 1 as expressed for statistical evaluation.

#### Sample group specificity

Hypergeometric tests followed by Bonferroni adjustment of *P* values were used to estimate statistical enrichment of gene expression in a particular sample group. Secondly, one-tailed Wilcoxon rank sum test was performed to compare the mean of a sample group expression to other groups to test whether gene is expressed at higher level. *P* value was corrected using Bonferroni if multiple groups were compared. Thirdly, fold change was computed between the tested groups.

### Development of immunological scores

#### Cytolytic score

To find genes most specifically expressed in CD8+ T cells/NK cells, specificity to these cell types was evaluated with the sample group specificity tests as described above (hypergeometric test adjusted *P* value < 1e-5, fold change > 1.5, Wilcoxon rank sum test adjusted *P* value < 0.01) using Hemap samples. Genes with significantly higher expression compared to B cells, HSC, erythroid cells, macrophage, monocyte, and dendritic cells were kept, resulting in 46 CD8+/NK cell-specific marker genes (Figure S1B). Known CD8+ T cell/NK cell-specific genes *GZMA, GZMB, PRF1, GNLY, GZMH*, and *GZMM* were chosen for further exploration, as they are directly related to cytolytic activity of T/NK cells.

Sorted BCL and AML samples and cell lines from all cancers, excluding T cell-like TCL lymphomas (because of transcriptomic similarity of TCL cancer cells and T/NK cells), were used to check whether these genes are expressed in pure cancer cell populations by requiring that expression values from pure samples (probeset noise distribution) were well separated from T/NK cells with high expression of the genes. As a second criteria, unsorted BCL and AML populations were compared to sorted BCL and AML samples to inspect if unsorted populations have samples with higher expression of the gene, indicating that increased signal is coming from T/NK cells. *GZMB* was filtered out, as it was highly expressed in a subset of pure samples. Geometric mean of gene expression for *GZMA, PRF1, GNLY, GZMH, GZMM* was computed followed by log2 transformation to be used as a proxy of cytolytic activity in hematological cancers.

#### HLA scores

To find genes related to HLA II antigen presentation, normal non-APC cells including CD4/CD8/regulatory/gamma-delta T cells, NK cells, erythroid lineage cells, and neutrophils were compared to normal APC cells including DCs, B cells, and macrophages to find HLA II genes that are expressed highly in APC cells, but not in any non-APC cells. Wilcoxon rank sum test was computed with option “greater” to find genes with higher expression in APC cells. Adjusted *P* value cutoff 0.001 and fold change cutoff > 4 were set to find significant genes, resulting in 350 genes. Genes were further filtered by computing fold change between each individual APC and non-APC cells to ensure genes are expressed higher in each APC cell type (median fold change > 2 to non-APC) resulting in total of 66 genes (Figure S4A and Table S4). Pairwise Spearman correlation of the genes overexpressed in APC was used to identify HLA II genes whose expression most highly correlated with each other using Hemap cancer samples (Figure S4B). The geometric mean of the HLA II genes *HLA-DMA, HLA-DMB, HLA-DPA1, HLA-DPB1, HLA-DRA*, and *HLA-DRB1* was defined as HLA II score.

Due to the ubiquitous expression of HLA I on all cell types, no filtering was necessary and the geometric mean of known HLA I genes *B2M, HLA-A, HLA-B*, and *HLA-C* was used to detect HLA I expression in the samples.

### Validation of immunological scores

#### Flow cytometry of AML patient samples

To analyze T and NK cell percentages in 37 AML BM samples, fresh diagnostic-phase BM aspirates in EDTA tubes were used. Antibodies according to Tables S1 and S4 were added to 50 μl of the BM sample, mixed and incubated for 15 min, washed with PBS + 0,1% NaN3, centrifuged, and the supernatant was discarded. Erythrocytes were lysed by incubating the sample in FACS Lysing Solution (BD) and the sample was washed with PBS + 0,1% NaN3, centrifuged, and the supernatant was discarded. The sample was resuspended into 0.5 ml FACSFlow Sheath Fluid (BD) and 200,000 events were acquired with FACSCanto (BD Pharmingen). Data were analyzed using FlowJo (10.0.8r1). For quantification of T/NK cells for cytolytic score validation, cell debris was excluded based on low forward scatter (FSC), lymphocytes were identified as CD45highSSClow cells, and T cells gated as CD3+ and NK cells as CD3-CD2+ out of lymphocytes (Figure S1D). Percentage of the sum of T and NK cells was calculated out of all non-debris cells. For quantification of HLA-DR+ AML blasts for HLA II score validation, cell debris was excluded based on low forward scatter (FSC), blasts were identified based on CD45 and SSC, and HLA-DR+ blasts gated as shown in Figure S4C.

#### RNA sequencing of AML patient samples for flow cytometry comparison

RNA sequencing was performed from the same 37 AML patient samples. Briefly, total RNA (2.5-5 μg) was extracted from BM mononuclear cells obtained by Ficoll-Paque gradient centrifugation using the miRNeasy kit (Qiagen) and depleted of ribosomal-RNA (Ribo-Zero™ rRNA Removal Kit, Epicentre) after purification, then reverse transcribed to double stranded cDNA (SuperScript™ Double-Stranded cDNA Synthesis Kit, Thermo Fisher Scientific). Sequencing libraries were prepared with Illumina compatible Epicentre Nextera™ Technology and ScriptSeq v2™ Complete kit (Illumina) and were purified with SPRI beads (Agencourt AMPure XP, Beckman Coulter) and library QC was evaluated on High Sensitivity chips by Agilent Bioanalyzer (Agilent). Paired-end sequencing with 100 bp read length was performed using Illumina HiSeq 2000. The reads were preprocessed as described previously (Kumar et al., 2017). Briefly, Trimmomatic was used to correct read data for low quality, Illumina adapters, and short read-length. Filtered paired-end reads were aligned to the human genome (GRCh38) using STAR (Dobin et al., 2013) with the guidance of EnsEMBL v82 gene models. Default 2-pass per-sample parameters were used, except that the overhang on each side of the splice junctions was set to 99. The alignments were then sorted and PCR duplicates were marked using Picard, feature counts were computed using SubRead (Liao et al., 2013), feature counts were converted to expression estimates using Trimmed Mean of M-values (TMM) normalization (Robinson and Oshlack, 2010) in edgeR (Robinson et al., 2010), and lowly expressed genomic features with counts per million (CPM) value ≤ 1.00 were removed. Default parameters were used, with exception that reads were allowed to be assigned to overlapping genome features in the feature counting. The immunological scores were calculated from TMM values based on geometric mean of selected genes, as described above.

#### Validation samples from Hemap

We used GSE13314 MALT lymphoma pathologic data from Chng *et al.* (Chng et al., 2009) (Table 1) to validate cytolytic score in lymphoma samples. GSMids with MALT immunohistochemistry data scored by a pathologist was compared to cytolytic score using Spearman’s correlation.

#### Comparison to other immunological scores

CIBERSORT (Newman et al., 2015) was computed for Hemap dataset with parameters perm=100 and QN=F. Similarly, R package “MCPcounter” command MCPcounter.estimate was used to infer immunology scores in Hemap dataset. GSVA was used to compute enrichment of Bindea *et al.* gene sets (Bindea et al., 2013) for Hemap data using parameter tau=0.25. Cytolytic score was correlated to CIBERSORT, MCP-counter, and Bindea *et al.* gene set scores using Pearson’s correlation in Hemap data.

### Analysis of microenvironment genes correlated to cytolytic score

Spearman correlation between gene expression level and cytolytic score was computed in unsorted cancer samples. Expression in CTL/NK cells was compared to a particular unsorted cancer sample group based on fold change. To distinguish genes that are CTL/NK cell-expressed, genes with significantly differential expression in CTL/NK cells compared to a particular normal cell type/purified cancer sample group were identified using the sample group specificity tests were performed as described above (hypergeometric test adjusted *P* value < 1e-3, fold change > 1.5, Wilcoxon rank sum test adjusted *P* value < 1e-5) using Hemap samples.

### Multiplexed immunohistochemistry (mIHC)

#### General

TMA blocks were sliced in 3.5 μm sections on Superfrost objective slides. We used 0.1% Tween-20 diluted in 10 mM Tris-HCL buffered saline pH 7.4 as washing buffer.

#### Tissue preparation

Tissue deparaffinization and rehydration was performed in xylene and graded ethanol series. Then, heat-induced epitope retrieval (HIER) was carried out in 10 mM Tris-HCl – 1 mM EDTA buffer (pH 9) in +99°C for 20 min (PT Module; Thermo Fisher Scientific). Tissue peroxide quenching with 0.9% H2O2 for 15 min was followed by protein blocking with 10% normal goat serum (TBS-NGS) for 15 min.

#### Staining

Primary antibody diluted in protein blocking solution as in (Tables S2 and S5) and secondary anti-mouse or antirabbit horseradish peroxidase-conjugated (HRP) antibody (Immunologic) diluted 1:1 in washing buffer were applied for 1h45min and 45 min, respectively. Tyramide signal was amplified (TSA; PerkinElmer) for 10 min. Primary antibodies and HRP activity were inactivated with HIER, followed by peroxide and protein block steps as described above. The second primary antibody with its matching HRP-conjugated secondary antibody diluted 1:5 in washing buffer were added and TSA signal amplified. We repeated HIER, peroxide block and protein block and applied two additional primary antibodies immunized in different species overnight in +4°C. AlexaFluor647 and AlexaFluor750 fluorochrome-conjugated secondary antibodies (Thermo Fisher Scientific) diluted 1:150 in washing buffer (45 min) and 4’,6-diamidino-2-phenylindole counterstain (DAPI; Roche/Sigma-Aldrich) diluted 1:250 in TBS (15 min) were added. Last, ProLong Gold (Thermo Fisher Scientific) was used to mount slides.

#### Imaging

Fluorescent images were acquired with the AxioImager.Z2 (Zeiss) microscope equipped with Zeiss Plan-Apochromat 20x objective (NA 0.8). Scanned images were acquired and converted to JPEG2000 format (95% quality). For representative images shown in figures, image channels were recolored using Fiji (Schindelin et al., 2012; Schneider et al., 2012), brightness and contrast were adjusted using identical parameters for images acquired using the same antibody panel to maintain comparability and representative regions of the images were selected.

#### Image analysis

Unfocused images were eliminated from the analysis. Cell segmentation and intensity measurements were computed based on adaptive Otsu thresholding and gradient intracellular intensity of grayscaled DAPI staining with the image analysis platform CellProfiler 2.1.2 (Carpenter et al., 2006). Cores with fewer than 1500 cells were eliminated from analysis. Cutoffs for marker positivity were based on staining intensity patterns of pooled cells of all samples and were confirmed visually (Tables S2 and S5, ‘mIHC panel’). Counts of all cells and cells positive for marker combinations were averaged across multiple cores of each patient when available. Immune cells were quantified as proportion of positive cells to all cells (e.g. proportion of CD68+ cells to total core cell count).

### Gene set enrichment analysis

Gene set enrichment analysis was computed using the command line version of GSEA (Subramanian et al., 2005). A total of 1645 genesets from MsigDB V5 c2 category gene sets (BIOCARTA, KEGG, REACTOME, SA, SIG, ST), MsigDB HALLMARKS, version 4 of NCI NATURE Pathway Interaction Database, and WIKIPW (6.2015) were used for enrichment analysis. Immunological scores were used as a continuous phenotype to rank genes using Pearson correlation as metric for ranking. Sample permutation and multiple hypothesis testing correction was done to obtain FDR for each gene set. Gene sets were limited to contain between 5 to 500 genes per gene set. Single sample specified in the result tables. This procedure allowed us enrichment score was assigned to significant pathways to identify most relevant correlations and to filter out for visualization and for Bindea gene sets using GSVA correlations difficult to interpret. package 1.24.0 (Hänzelmann et al., 2013) in R.

### Genomic correlations with immunological features

#### Feature generation from multilevel data

Feature matrix generation, pairwise analysis run, and feature-specific filtering was based on the TCGA featurematrix pipeline available in https://github.com/cancerregulome/gidget/tree/master/commands/feature_matrix_construction. Continuous/discrete numeric data matrices were generated for the analyzed datasets Hemap, TCGA AML and DLBCL, Chapuy *et al.* DLBCL, and Glass *et al.* AML including clinical, genomic and immunologic features. In case of categorical features, binary indicator features were generated to compare i) each categorical feature to the rest of the samples, ii) two categorical features to rest of the samples and iii) comparing two categorical features against each other. Feature types (gene expression, protein expression, clinical, methylation, CNV, mutations, sample annotation) were distinguished from each other. Missing values were assigned as NA. To account for differential methylation within the same locus, probes associated with each gene were first correlated to gene expression using Spearman’s correlation (*P* < 0.05) and divided into positive and negatively correlated sets. Probes with standard deviation below 0.1 were removed. Mean methylation for these sets were computed to obtain two methylation features per gene, with positive and negative association to gene expression.

#### Feature statistical analysis

Spearman correlation was used to for numeric-numeric and numeric-binary feature pairs (in R use=”pairwise.complete.obs”), while one-tailed Fisher’s exact test test of co-occurrence was used for binary-binary feature analysis. To keep the number of comparisons smaller and statistically reliable, only features with at least 5 observations were used in the analysis.

For *P* value adjustment, the number of features and intrinsic correlation are very different for different data pairs. Therefore, we performed separate statistical tests: The first test was to find whether features of a gene (methylation, mutation, CNV, gene expression) are correlated with each other (to identify alterations of the gene itself associated with each other), followed by Benjamini-Hochberg (BH) correction of the obtained *P* values. The second test was to assess whether features of a gene are correlated to features of other genes (to identify e.g. driver alterations associated with immunological features) and included additional feature types, such as clinical variables and sample annotations, followed by BH correction of *P* values. Multiple hypothesis testing was performed separately for the correlation and Fisher’s test results, as these produce different *P* value distributions. Similarly, different significance level cutoffs were used for different data pairs. Methylation-methylation and CNV-CNV pairs were omitted. FDR cutoff was set to 0.1, except for mutations FDR < 0.25 was permitted. Further filtering criteria are

### Differential methylation analysis

#### ERRBS (Glass et al. AML)

GSE86952 raw aligned ERRBS AML methylation data were analyzed using the methylSig (Park et al., 2014) R package to identify methylation changes in patients with low HLA II expression or PML-RARA mutation compared to rest of the samples. Intersection of samples with both methylation, gene expression, and mutation data was 106 samples. Data files were read in R using the methylSigReadData command with parameters context=”CpG” and destranded=TRUE. methylSigCalc command was run to obtain differentially methylated CpGs with parameters min.per.group=5 to require a minimum of 5 CpGs per group for calling differential methylation. All significant DMCs with FDR < 0.05 and absolute methylation change > 25% for *CIITA* genomic locus were obtained.

#### Illumina 450k (TCGA)

For comparison of differentially methylated immune checkpoint genes between TCGA AML and DLBCL samples, beta values of Illumina 450k probes within 1 kb of the transcription start site of the genes were averaged to obtain a single value representing methylation of the gene promoter area. M values were calculated from mean beta values using log2(beta/(1-beta)) and differential methylation analysis was performed using limma. For the corresponding differential gene expression analysis, RNA-seq read counts were converted to cpm and normalized using limma voom with quantile normalization, and differential expression analysis was performed using limma.

*CCLE* For correlation of gene expression to methylation in CCLE RRBS data, CpG and TSS 1 kb methylation values of each gene were averaged and the resulting mean methylation value was compared with gene expression using Spearman correlation.

### Cell culture experiments

For drug treatment experiments, MOLM13 cells were plated on flat-bottom 96-well plates at 50,000 cells/well in a volume of 100 μL. Cell were treated with 10, 100 or 1,000 nM decitabine (Selleck) or DMSO as a control, both in the presence or absence of 10 ng/mL recombinant human interferon gamma (Peprotech), all conditions in triplicate wells. After 3 days, 25 μL of cell suspension from each well was washed with 100 μL PBS-EDTA and stained with 5 μL of HLA-DR-FITC (clone G46-6, BD BioSciences) or isotype control (clone G155-178, BD BioSciences) antibodies or left unstained in a total volume of 25 μL PBS-EDTA and antibodies. Cells were then washed with 100 μL PBS-EDTA, resuspended to 50 μL PBS-EDTA, and 10,000 events were acquired on a FACSVerse flow cytometer (BD BioSciences). Flow cytometry data were analyzed using FlowJo 10.0.8r1. Viable cells were gated using forward (FSC-A) and side scatter (SSC-A), followed by gating for singlets using FSC-A and FSC-H. HLA-DR+ cells were gated such that untreated isotype control-stained cells were gated negative. Final HLA-DR+ cell percentages were obtained by subtracting isotype control-stained HLA-DR+ cell percentages (mean of triplicate wells) from HLA-DR Hemap to make sure findings are not due to differences antibody-stained HLA-DR+ cell percentages at each in survival cohorts. treatment condition.

### Immune checkpoint gene list curation

A list of co-stimulatory and co-inhibitory ligands and immunomodulatory enzymes known to be expressed on APCs or T/NK cell target cells was curated based on the literature. T cell ligands were based largely on a comprehensive review on T cell co-stimulation (Chen and Flies, 2013), and the list was supplemented with NK cell receptor ligands and immunomodulatory enzymes. References for the receptor-ligand interactions and their stimulatory or inhibitory effect on T/NK cells are listed in Table S5.

### Antigen expression analysis

Genes expressed only in normal or cancer cells were identified using the sample group specificity tests were performed as described above (hypergeometric test adjusted *P* value < 1e-2, fold change > 1.25, Wilcoxon rank sum test adjusted *P* value < 1e-5) using Hemap samples. Genes expressed in cancer were required to be expressed highly in > 5% of the patients in at least one disease based on mixture model categories (high vs. low expressed/not detected) and not expressed in normal cells. GTEx database (GTEx Consortium, 2015) V6 RNA-seq gene median RPKM values for each tissue were used to find genes specific to testis when compared to other tissues (excluding ovary). Testis genes were defined to have < 0.25 RPKM expression in all other tissues, resulting in a total of 1,563 genes common between Hemap and GTEx datasets. This list of genes was filtered to contain only coding genes expressed in Hemap cancer samples, resulting in 59 genes. CCLE cell line data for hematological cancers was also used to filter out genes not expressed > 0.5 RPKM levels in at least 5 cell lines, resulting in a final antigen genelist containing 27 genes. Number of expressed CGA genes was computed for each sample using mixture model-discretized gene expression values for Hemap data, and using a cutoff of 0.25 RPKM for CCLE data. For Chapuy *et al.* DLBCL, patient LS2208 with testicular DLBCL was omitted from the analysis.

### Survival analysis

#### Univariate analysis

Cox regression available in R package ‘survival’ version 2.42-4 was applied for univariate analyses of numeric immunology scores, including HLA I score, HLA II score, and CGA number in myeloma, AML, and DLBCL datasets, and cytolytic score in AML and DLBCL where unsorted samples were available (for myeloma GSE19784, GSE16716, GSE24080, for DLBCL GSE10846, GSE11318, and GSE17372 and for AML GSE10358, GSE12662, GSE12417, GSE14468, GSE6891). Furthermore, all co-stimulatory genes and individual CGAs were included in the analysis. Additionally, non-T/NK expressed genes with expression fold change > 2 between CTLs/NK cells and the unsorted cancer samples and correlation with cytolytic score above 0.4 (as in Figures 2 and S2). Well-known prognostic markers, including ISS for myeloma and IPI for DLBCL were also included in the analysis. Survival data were analyzed also for each individual study in

#### Multivariate analysis

Features correlated to overall survival in univariate analysis for each disease were selected for multiple regression analyses computed in Hemap myeloma and DLBCL datasets to evaluate their prognostic significance. Features were filtered using an adjusted *P* value cutoff 0.2 to reduce the number of features for the multiple regression analysis and to decrease the the false discovery rate. Regularized Cox regression model available in glmnet 2.0.16 R package was used to fit the Cox model. L1 and L2 norm ratio (alpha parameter) was optimized using 10-fold cross-validation and alpha values from 0 to 1 with 0.05 increments. To reduce variability in the model, cross-validation for each alpha value was iterated 100 times. Alpha and lambda with the lowest mean fitting error were used for the final model fitting. Independent test datasets were used for model validation. Hemap RCHOP-treated samples from GSE10846 and GSE17372 were used for training and GSE98588 for testing in DLBCL, and for myeloma GSE19874 was used for model training and GSE24080 for model testing. Prognostic index (PI) was computed for each sample as in (Royston and Altman, 2013) for training and test sets. Cox proportional hazards model and Kaplan-Meier plots were used to compare model performance between training and test sets. PI with a similar hazard ratio and a low overall *P* value were used to verify that a set of distinct immunology features could be used to distinguish different patient outcomes independently of the dataset where the model was generated.

### Data visualization

R package “ComplexHeatmap” (Gu et al., 2016) was used for drawing heatmaps and oncoprints and “ggplot2” for drawing boxplots, barplots, and dot plots. Gene expression Z-scores were used for t-SNE map visualization to denote samples with low and high expression (low: < −2 to −1 and high 1 to > 2). For Hemap dataset, e-staining was used for gene expression visualization for mixture model components (not detected, low, and high) as described above.

### Statistical analysis

The statistical details of all experiments are reported in the text, figure legends, and figures, including statistical analysis performed, statistical significance, and counts. Significance codes correspond to *P* values or FDR as follows: * < 0.05, ** < 0.01, *** < 0.001, **** <0.0001. In boxplots, horizontal line indicates the median, boxes indicate the interquartile range, and whiskers extend from the hinge to the smallest/largest value at most 1.5 * IQR of the hinge. Generally, nonparametric methods including Spearman correlation and Wilcoxon rank sum test were used for statistical analyses.

### Data and software availability

Software used for the analyses are described and referenced in the individual Method Details subsections and are listed in the Key Resources Table. Scripts used to generate results are available upon request. Hemap data can be queried and visualized from http://hemap.uta.fi.

## REAGENTS AND RESOURCES

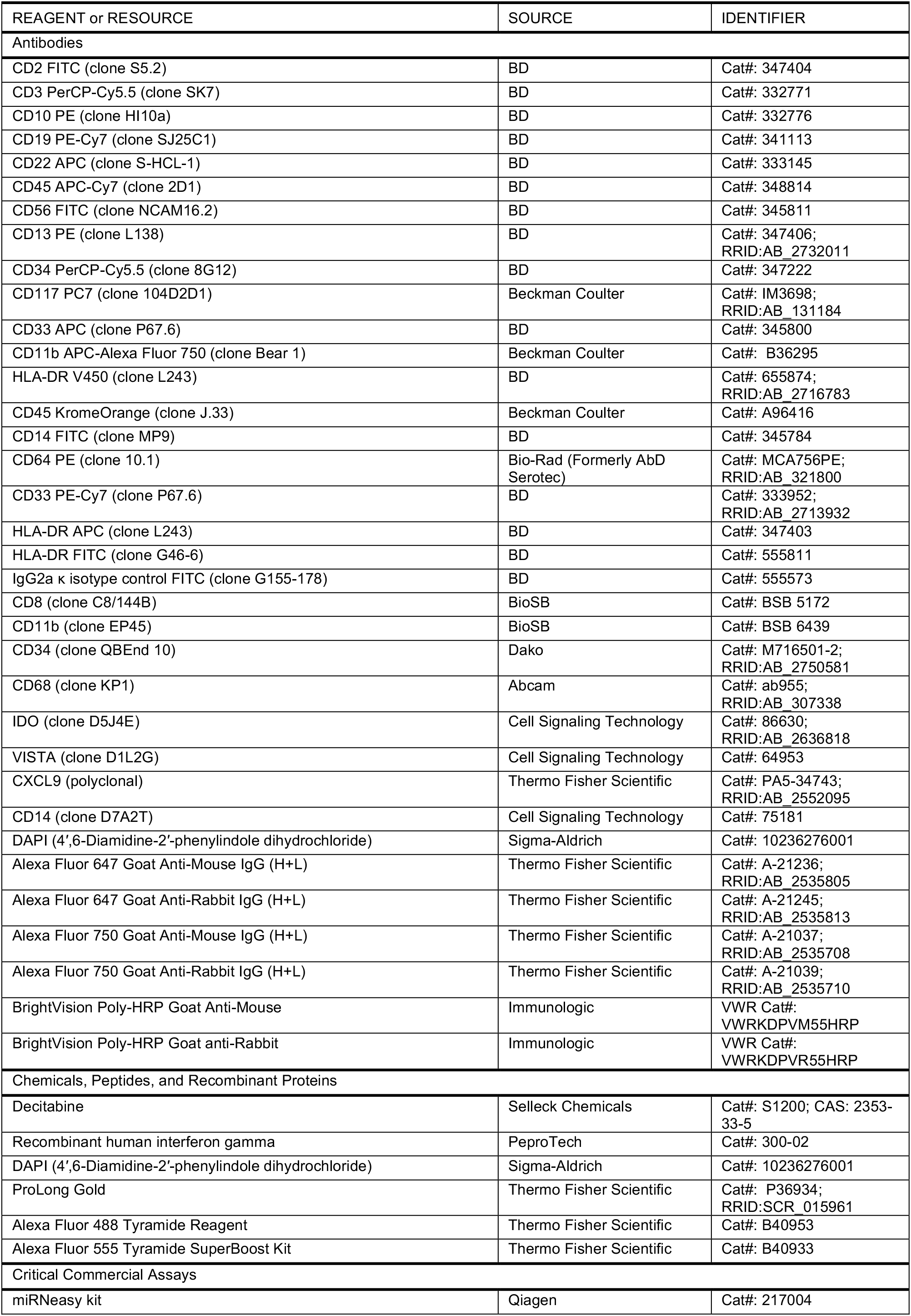
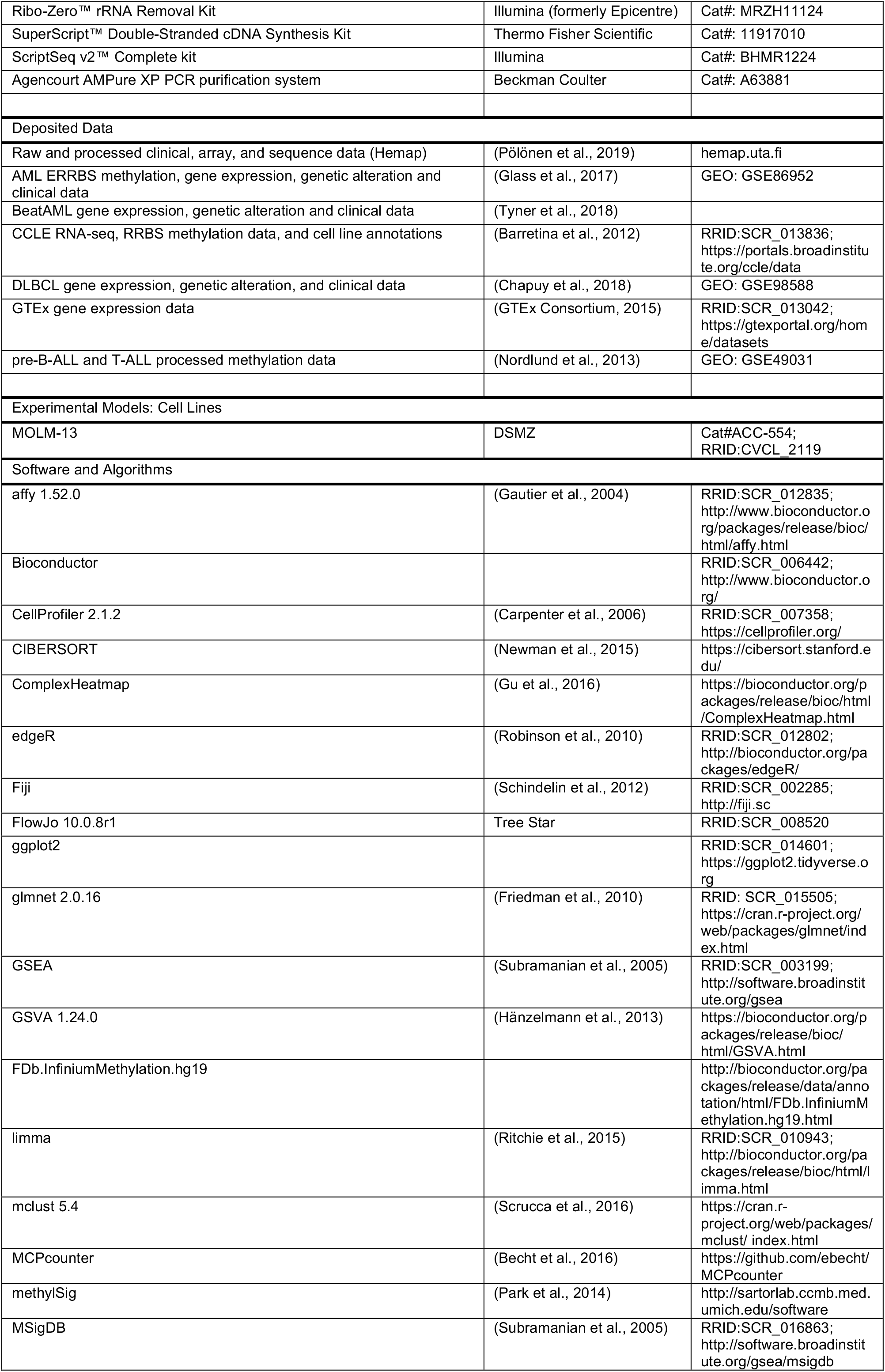
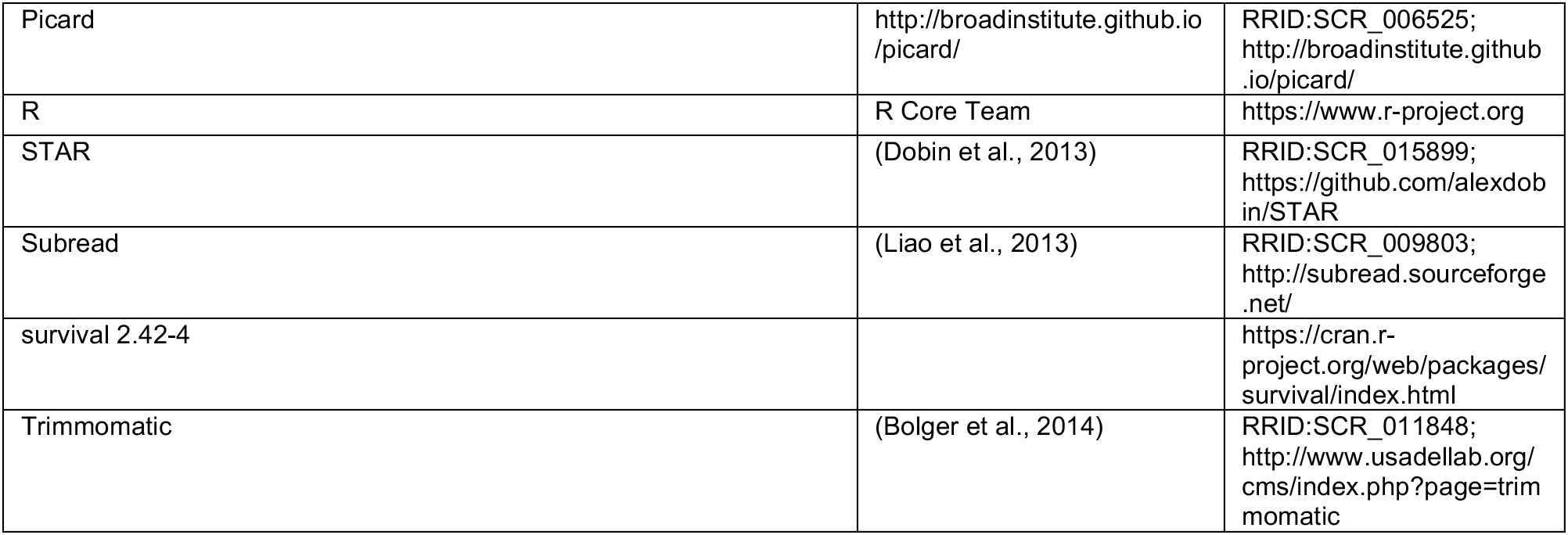

### ACKNOWLEDGMENTS

We thank the Helsinki Biobank and the Digital and Molecular Pathology Unit supported by Helsinki University and Biocenter Finland for digital microscopy services. We acknowledge staff at the Institute of Molecular Medicine Finland (FIMM), including the FIMM sequencing lab and FIMM bioinformatics unit for preparing the AML RNA-Seq validation data, and the High Throughput Biomedicine unit. High Throughput Biomedicine. We thank the personnel of Hematology Research Unit Helsinki for their excellent technical assistance.

The results published here are in part based upon data generated by The Cancer Genome Atlas project (dbGaP Study Accession: phs000178.v8.p7) established by the NCI and NHGRI. Information about TCGA and the investigators and institutions who constitute the TCGA research network can be found at http://cancergenome.nih.gov. The Genotype-Tissue Expression (GTEx) Project was supported by the Common Fund of the Office of the Director of the National Institutes of Health, and by NCI, NHGRI, NHLBI, NIDA, NIMH, and NINDS.

This study was supported by the Finnish Cancer Organizations, Sigrid Juselius Foundation, Relander Foundation, State funding for university-level health research in Finland, and HiLife fellow funds from the University of Helsinki.

### AUTHOR CONTRIBUTIONS

O.D., P.P, M.K., M.H. and S.M. conceived the study. O.D. and P.P. designed and performed the analyses. M.K. provided input on the study design and interpreted the data. O.B. performed multiplexed immunohistochemistry. J.M. developed data analysis tools. A.K. performed AML RNA-seq data analysis and C.H. supervised sample collection for RNA-seq. S.S. generated the flow cytometry data. S.-K.L., L.M., and S.L. collected and constructed the DLBCL TMA. M.N. and O.L. supervised Hemap data collection and harmonization. O.D. and P.P. wrote the manuscript with input from S.M., M.H., and M.K. and contributions from all authors. S.M. and M.H. supervised the project.

