## Supplementary Materials for "Immunogenomic landscape of hematological malignancies"

#### SUPPLEMENTAL FIGURES

Figure S1

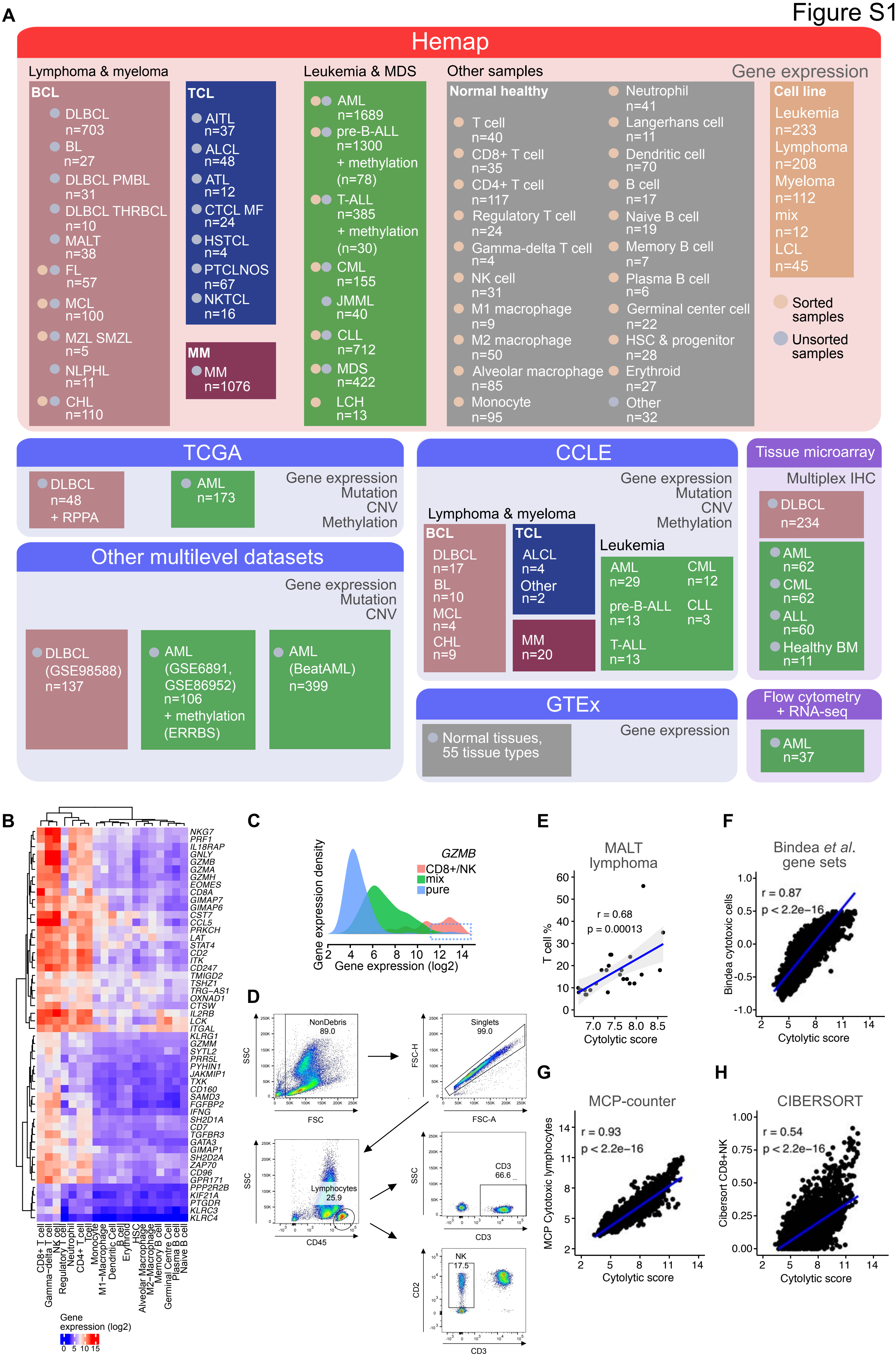
A
Hemap
Lymphoma & myeloma
BCL
DLBCL n=703
BL n=27
DLBCL PMBL n=31
DLBCL THRBCL n=10
MALT n=38
FL n=57
MCL n=100
MZL SMZL n=5
NLPHL n=11
CHL n=110
TCL
AITL n=37
ALCL n=48
ATL n=12
CTCL MF n=24
HSTCL n=4
PTCLNOS n=67
NKTCL n=16
MM
MM n=1076
Leukemia & MDS
AML n=1689
pre-B-ALL n=1300 + methylation (n=78)
T-ALL n=385 + methylation (n=30)
CML n=155
JMML n=40
CLL n=712
MDS n=422
LCH n=13
Other samples
Normal healthy
T cell n=40
CD8+ T cell n=35
CD4+ T cell n=117
Regulatory T cell n=24
Gamma-delta T cell n=4
NK cell n=31
M1 macrophage n=9
M2 macrophage n=50
Alveolar macrophage n=85
Monocyte n=95
Neutrophil n=41
Langerhans cell n=11
Dendritic cell n=70
B cell n=17
Naive B cell n=19
Memory B cell n=7
Plasma B cell n=6
Germinal center cell n=22
HSC & progenitor n=28
Erythroid n=27
Other n=32
Gene expression
Cell line
Leukemia n=233
Lymphoma n=208
Myeloma n=112
mix n=12
LCL n=45
Sorted samples
Unsorted samples
TCGA
DLBCL n=48 + RPPA
AML n=173
Gene expression
Mutation
CNV
Methylation
CCLE
Gene expression
Mutation
CNV
Methylation
Lymphoma & myeloma
BCL
DLBCL n=17
BL n=10
MCL n=4
CHL n=9
TCL
ALCL n=4
Other n=2
MM n=20
Leukemia
AML n=29
CML n=12
pre-B-ALL n=13
CLL n=3
T-ALL n=13
Tissue microarray
Multiplex IHC
DLBCL n=234
AML n=62
CML n=62
ALL n=60
Healthy BM n=11
Other multilevel datasets
Gene expression
Mutation
CNV
DLBCL (GSE98588) n=137
AML (GSE6891, GSE86952) n=106 + methylation (ERRBS)
AML (BeatAML) n=399
GTEx
Normal tissues, 55 tissue types
Gene expression
Flow cytometry + RNA-seq
AML n=37
B
NKG7
PRF1
IL18RAP
GNLY
GZMB
GZMA
GZMH
EOMES
CD8A
GIMAP7
GIMAP6
CST7
CCL5
PRKCH
LAT
STAT4
CD2
ITK
CD247
TMIGD2
TSHZ1
TRG−AS1
OXNAD1
CTSW
IL2RB
LCK
ITGAL
KLRG1
GZMM
SYTL2
PRR5L
PYHIN1
JAKMIP1
TXK
CD160
SAMD3
FGFBP2
IFNG
SH2D1A
CD7
TGFBR3
GATA3
GIMAP1
SH2D2A
ZAP70
CD96
GPR171
PPP2R2B
KIF21A
PTGDR
KLRC3
KLRC4
CD8+ T cell
Gamma-delta T cell
NK cell
Regulatory T cell
Neutrophil
CD4+ T cell
Tcell
Monocyte
M1−Macrophage
Dendritic Cell
B cell
Erythroid
HSC
Alveolar Macrophage
M2−Macrophage
Memory B cell
Germinal Centre Cell
Plasma B cell
Naive B cell
Gene expression (log2)
0 5 10 15
C
GZMB
CD8+/NK
mix
pure
Gene expression density
Gene expression (log2)
2 4 6 8 10 12 14
D
NonDebris 89.0
SSC
FSC
Singlets 99.0
FSC-H
FSC-A
Lymphocytes 25.9
CD45
CD3 66.6
CD3
NK 17.5
CD2
E
MALT lymphoma
r = 0.68
p = 0.00013
T cell %
Cytolytic score
F
Bindea et al. gene sets
r = 0.87
p < 2.2e−16
Bindea cytotoxic cells
Cytolytic score
G
MCP-counter
r = 0.93
p < 2.2e−16
MCP Cytotoxic lymphocytes
Cytolytic score
H
CIBERSORT
r = 0.54
p < 2.2e−16
Cibersort CD8+NK
Cytolytic score

**Figure S1. Schematic of samples included in the study and assessment of cytotoxic lymphocyte infiltration in hematological malignancies using gene expression data. Related to Figure 1. A.** Schematic of all the datasets, sample numbers, and data types used in the analyses. **B.** Median expression of genes most enriched in normal CTLs and NK cells shown as a heatmap. Rows and columns are clustered using euclidean distance and complete linkage. **C.** Density plot of *GZMB* gene expression in purified cancer cells, unsorted cancer samples and CD8+T cell/NK cell populations, showing a subset of purified samples highly expressing *GZMB*. **D.** Flow cytometry gating strategy to identify percentages of T cells (CD3+) and NK cells for comparison with RNA-seq data (related to Figure 1C). **E.** Cytolytic score validation in Hemap MALT lymphoma. Correlation (Spearman) between cytolytic score and reported T cell percentage based on pathological analysis. **F-H.** Scatterplots comparing cytolytic score and Bindea *et al.* "Cytotoxic cells" gene set score, MCP-counter "Cytotoxic lymphocytes", and CIBERSORT sum of "T cells CD8", "NK cells resting", and "NK cells activated" fractions. Spearman correlation between each comparison is shown.

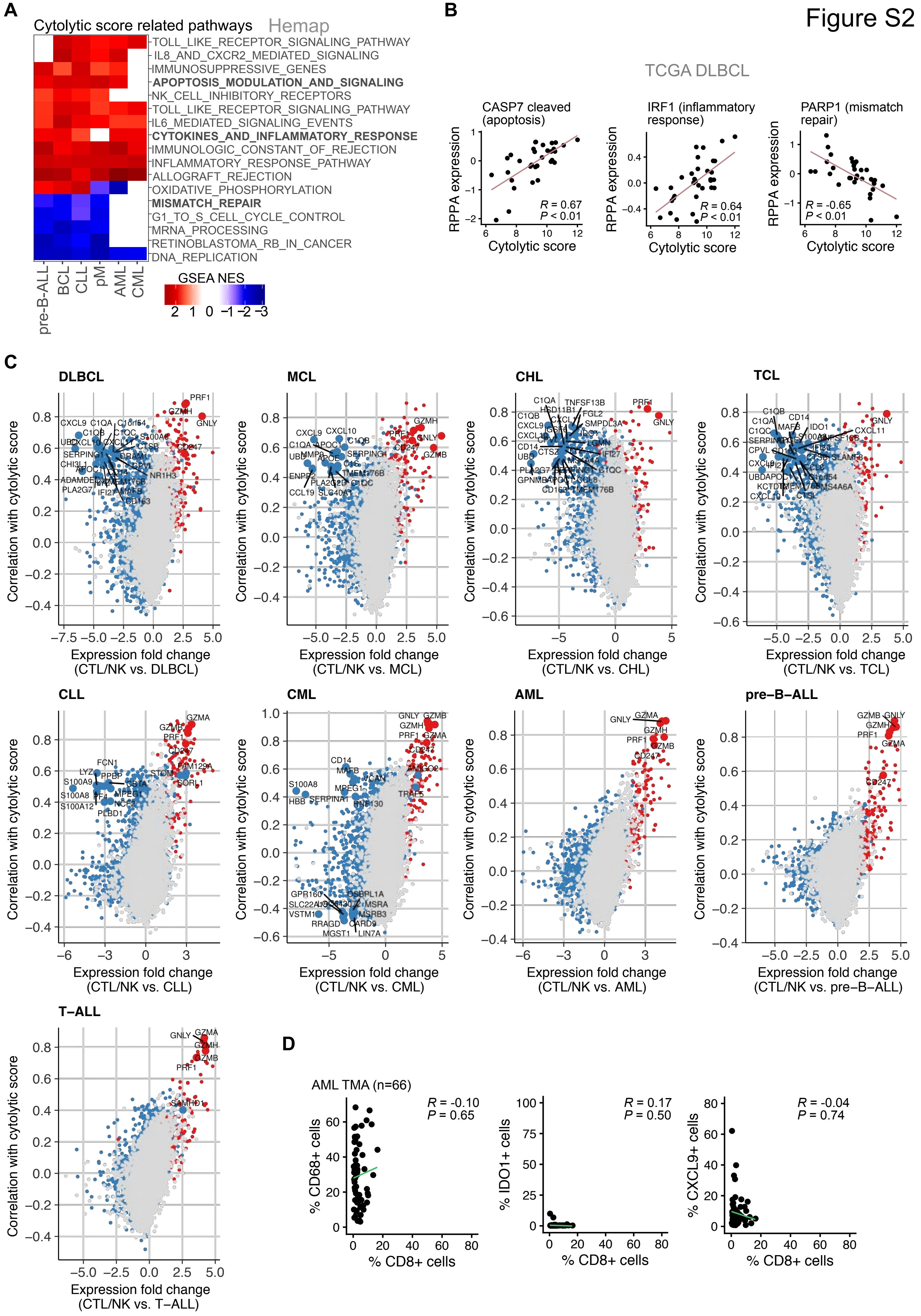
Figure S2
A
Cytolytic score related pathways
Hemap
TOLL_LIKE_RECEPTOR_SIGNALING_PATHWAY
IL8_AND_CXCR2_MEDIATED_SIGNALING
IMMUNOSUPPRESSIVE_GENES
APOPTOSIS_MODULATION_AND_SIGNALING
NK_CELL_INHIBITORY_RECEPTORS
TOLL_LIKE_RECEPTOR_SIGNALING_PATHWAY
IL6_MEDIATED_SIGNALING_EVENTS
CYTOKINES_AND_INFLAMMATORY_RESPONSE
IMMUNOLOGIC_CONSTANT_OF_REJECTION
INFLAMMATORY_RESPONSE_PATHWAY
ALLOGRAFT_REJECTION
OXIDATIVE_PHOSPHORYLATION
MISMATCH_REPAIR
G1_TO_S_CELL_CYCLE_CONTROL
MRNA_PROCESSING
RETINOBLASTOMA_RB_IN_CANCER
DNA_REPLICATION
pre-B-ALL
BCL
CLL
pM
AML
CML
GSEA NES
2 1 0 −1 −2 −3
B
TCGA DLBCL
CASP7 cleaved (apoptosis)
R = 0.67
P < 0.01
IRF1 (inflammatory response)
R = 0.64
P < 0.01
PARP1 (mismatch repair)
R = -0.65
P < 0.01
RPPA expression
Cytolytic score
C
DLBCL
MCL
CHL
TCL
CLL
CML
AML
pre−B−ALL
T−ALL
Correlation with cytolytic score
Expression fold change (CTL/NK vs. DLBCL)
Expression fold change (CTL/NK vs. MCL)
Expression fold change (CTL/NK vs. CHL)
Expression fold change (CTL/NK vs. TCL)
Expression fold change (CTL/NK vs. CLL)
Expression fold change (CTL/NK vs. CML)
Expression fold change (CTL/NK vs. AML)
Expression fold change (CTL/NK vs. pre−B−ALL)
Expression fold change (CTL/NK vs. T−ALL)
PRF1
GZMH
GNLY
GZMA
GZMB
CD247
D
AML TMA (n=66)
R = -0.10
P = 0.65
% CD68+ cells
R = 0.17
P = 0.50
% IDO1+ cells
R = -0.04
P = 0.74
% CXCL9+ cells
% CD8+ cells

**Figure S2. Gene expression correlations with cytolytic score across hematological malignancies. Related to Figure 2. A.** Heatmap of gene set enrichment analysis (GSEA) normalized enrichment scores (NES) for significantly enriched pathways correlated with cytolytic activity in each Hemap cancer type. Pathways related to D highlighted. **B.** TCGA DLBCL RPPA protein expression correlations with cytolytic score to validate pathway enrichment profiles highlighted in C for DLBCL. **C.** Scatter plots of genes correlated with cytolytic score in each cancer types as in Figure 2A. **D.** Scatter plots comparing the percentage of CD8+ cells (CTLs) out of total cells with the percentages of CD68+ cells (macrophages), IDO1+ cells, and CXCL9+ cells out of total cells in the AML IHC cohort (n = 62). Spearman correlation coefficients and *P* values adjusted using the BH method are shown.

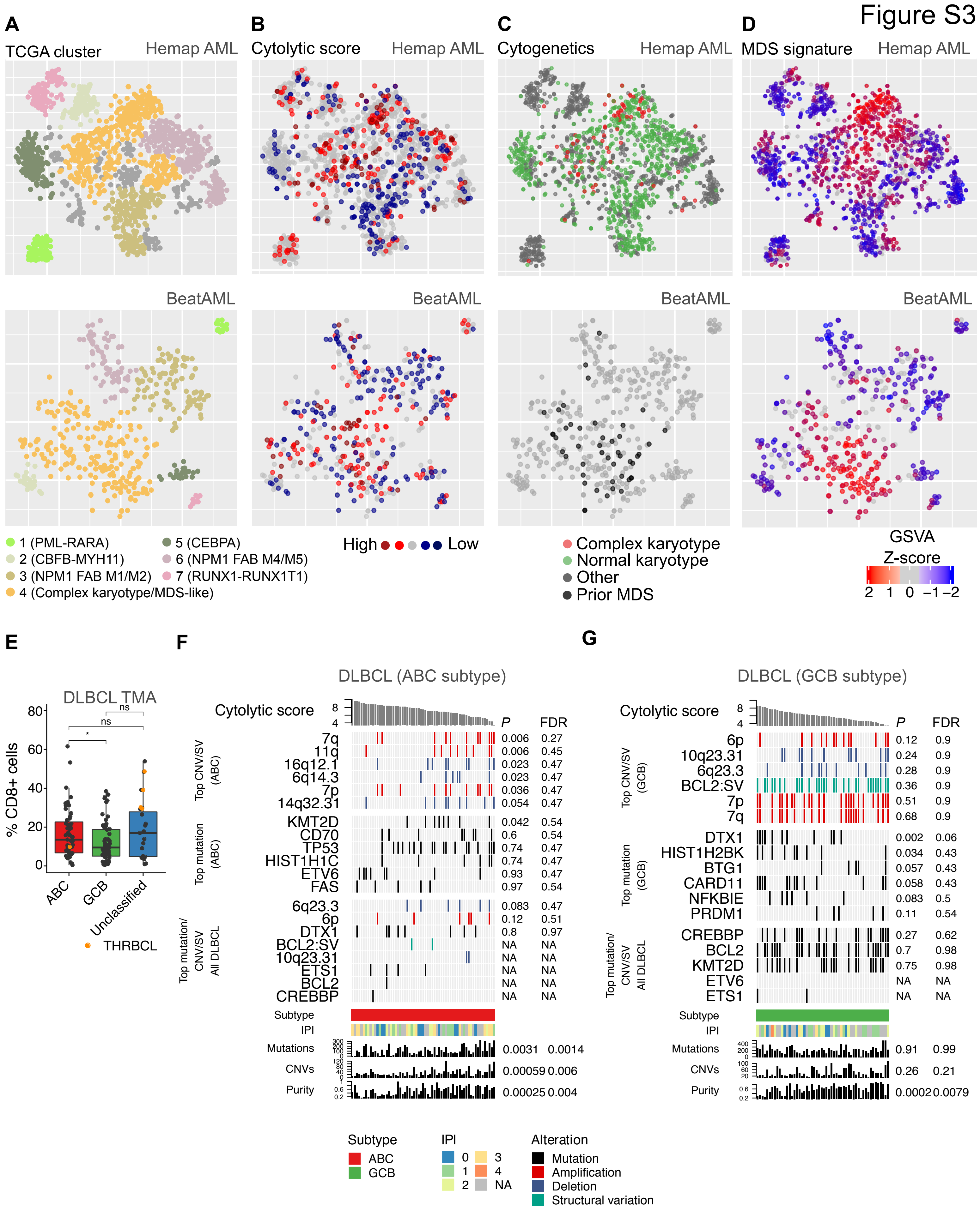

Figure S3
A
TCGA cluster
Hemap AML
BeatAML
1 (PML-RARA)
2 (CBFB-MYH11)
3 (NPM1 FAB M1/M2)
4 (Complex karyotype/MDS-like)
5 (CEBPA)
6 (NPM1 FAB M4/M5)
7 (RUNX1-RUNX1T1)
B
Cytolytic score
High
Low
C
Cytogenetics
Complex karyotype
Normal karyotype
Other
Prior MDS
D
MDS signature
GSVA Z-score
E
DLBCL TMA
% CD8+ cells
ABC
GCB
Unclassified
THRBCL
ns
*
F
DLBCL (ABC subtype)
Cytolytic score
Top CNV/SV (ABC)
Top mutation (ABC)
Top mutation/CNV/SV All DLBCL
P
FDR
7q 0.006 0.27
11q 0.006 0.45
16q12.1 0.023 0.47
6q14.3 0.023 0.47
7p 0.036 0.47
14q32.31 0.054 0.47
KMT2D 0.042 0.54
CD70 0.6 0.54
TP53 0.74 0.47
HIST1H1C 0.74 0.47
ETV6 0.93 0.47
FAS 0.97 0.54
6q23.3 0.083 0.47
6p 0.12 0.51
DTX1 0.8 0.97
BCL2:SV NA NA
10q23.31 NA NA
ETS1 NA NA
BCL2 NA NA
CREBBP NA NA
Subtype
IPI
Mutations 0.0031 0.0014
CNVs 0.00059 0.006
Purity 0.00025 0.004
G
DLBCL (GCB subtype)
Cytolytic score
Top CNV/SV (GCB)
Top mutation (GCB)
Top mutation/CNV/SV All DLBCL
P
FDR
6p 0.12 0.9
10q23.31 0.24 0.9
6q23.3 0.28 0.9
BCL2:SV 0.36 0.9
7p 0.51 0.9
7q 0.68 0.9
DTX1 0.002 0.06
HIST1H2BK 0.034 0.43
BTG1 0.057 0.43
CARD11 0.058 0.43
NFKBIE 0.083 0.5
PRDM1 0.11 0.54
CREBBP 0.27 0.62
BCL2 0.7 0.98
KMT2D 0.75 0.98
ETV6 NA NA
ETS1 NA NA
Subtype
IPI
Mutations 0.91 0.99
CNVs 0.26 0.21
Purity 0.0002 0.0079
Subtype
ABC
GCB
IPI
0 1 2 3 4 NA
Alteration
Mutation
Amplification
Deletion
Structural variation

**Figure S3. Genetic alterations and molecular subtypes associated with cytolytic score. Related to Figure 3. A-D.** Visualization of Hemap AML and BeatAML samples using a t-SNE representation. (A) Clusters, (B) cytolytic score, (C) cytogenetics in Hemap AML and MDS status in BeatAML, and (D) MDS signature are colored on the t-SNE maps. Key characteristics of the clusters are annotated. **E.** Comparison of CD8+ cell percentages out of all cells between DLBCL molecular subtypes based on quantitative multiplex immunohistochemistry performed on tissue microarrays. ABC and GCB subtypes are compared using a one-tailed Wilcoxon rank sum test. THRBCL cases are colored in yellow. **F.** Genetic alterations associated with cytolytic score in DLBCL patients of ABC subtype (n = 63, GSE98588) shown as an oncoprint where columns corresponding to a sample are ranked by cytolytic score and mutations, copy number variations (CNVs), structural variations (SVs), cell of origin, IPI risk score, mutation burden, CNV number, and sample purity are plotted on rows. Discrete state/class is indicated as color (genetic aberrations and sample categories) and continuous values are represented as barcharts (score or percentage values). *P* values and FDR for correlations between cytolytic score and genetic aberrations are shown. Top CNVs/SVs correlated with cytolytic score in ABC are shown at the top, top mutations correlated with cytolytic score in ABC in the middle, and mutations and CNVs correlated with cytolytic score in all DLBCLs from Figure 3E are shown at the bottom. **G.** Genetic alterations associated with cytolytic score in DLBCL patients of GCB subtype (n = 54, GSE98588) are shown as in C.

### Figure S4

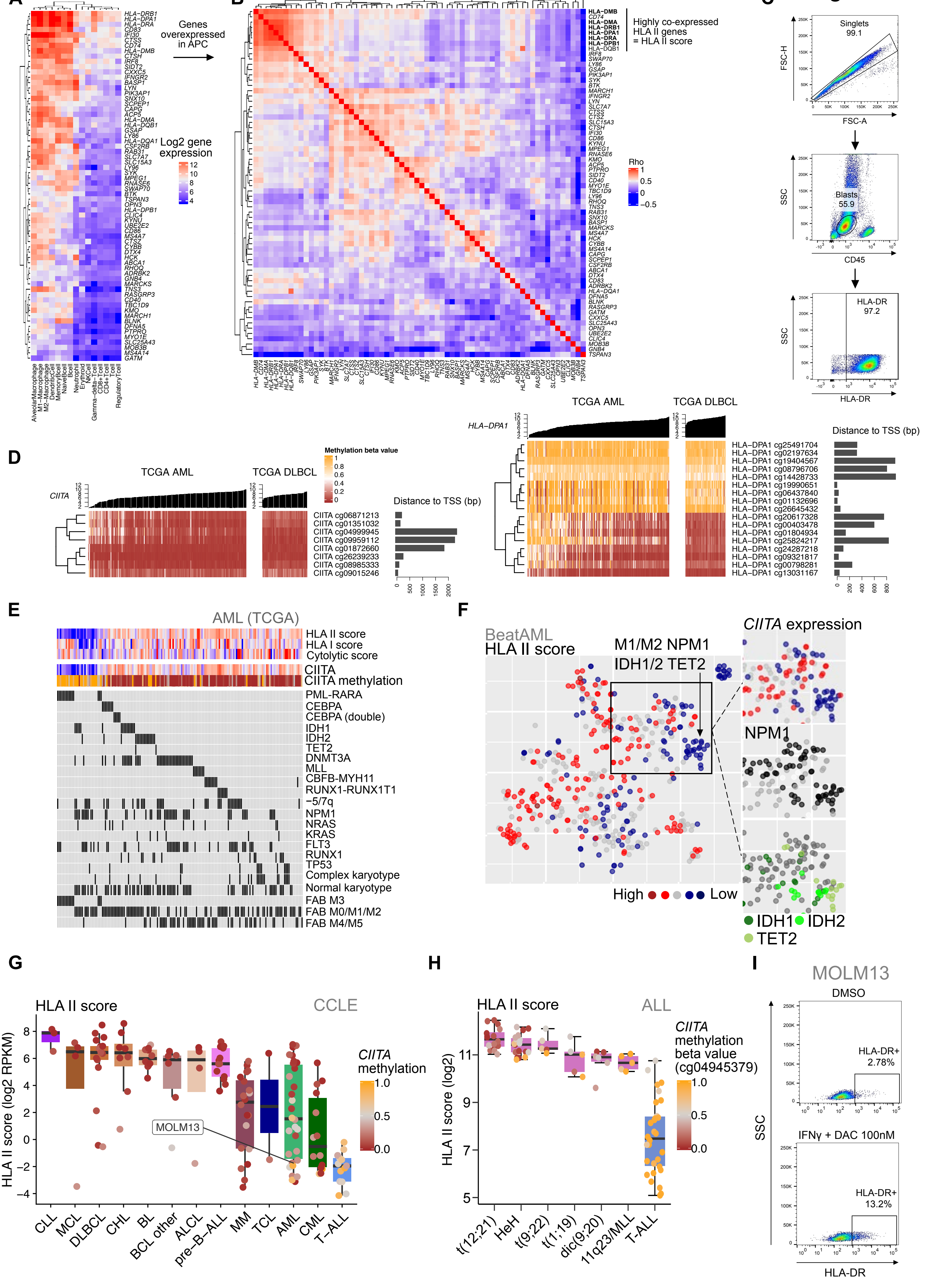

A
APC
Non-APC
Genes overexpressed in APC
Log2 gene expression
B
Highly co-expressed HLA II genes = HLA II score
Rho
C
Singlets 99.1
FSC-H
FSC-A
SSC
Blasts 55.9
CD45
HLA-DR 97.2
HLA-DR
D
TCGA AML
TCGA DLBCL
Methylation beta value
Distance to TSS (bp)
E
AML (TCGA)
HLA II score
HLA I score
Cytolytic score
CIITA
CIITA methylation
F
BeatAML
HLA II score
M1/M2 NPM1
IDH1/2 TET2
CIITA expression
NPM1
High
Low
IDH1
IDH2
TET2
G
HLA II score
CCLE
HLA II score (log2 RPKM)
CIITA methylation
MOLM13
H
HLA II score
ALL
HLA II score (log2)
CIITA methylation beta value (cg04945379)
I
MOLM13
DMSO
HLA-DR+ 2.78%
IFNγ + DAC 100nM
HLA-DR+ 13.2%
SSC
HLA-DR

**Figure S4. Genetic and epigenetic alterations associated with HLA gene expression. Related to Figure 4. A.** Heatmap of log2 mean expression of genes significantly (FDR < 0.01 and FC > 2) differentially expressed between APC and non-APC normal cell types. Rows and columns are clustered using Euclidean distance and complete linkage. **B.** Pairwise gene correlations (Spearman) for genes from A for Hemap tumor samples are shown as a correlation matrix heatmap. HLA II genes included in the HLA II score based on their high correlation with each other are indicated in bold. **C.** Flow cytometry gating strategy to identify percentages of HLA-DR+ blasts in diagnostic AML BM sample for comparison with RNA-seq data (related to Figure 4D). **D.** *CIITA* and *HLA-DPA1* methylation values are shown for each probe associated with the gene for TCGA AML and DLBCL. Samples are ranked by gene expression indicated above the heatmap. Probe distances (bp) from transcription start site (TSS) are shown as barplot on the right. **E.** Oncoprint of TCGA AML genetic alterations organized by HLA II score. Immunological scores, *CIITA* expression and methylation, karyotype, common genetic alterations, and morphological FAB type are shown. **F.** HLA II score colored on the BeatAML t-SNE map. CIITA expression, and *NPM1* and *IDH1*, *IDH2,* and *TET2* mutation status is labeled for cluster with low HLA II score. **G.** HLA II score of CCLE hematological cell lines. Dot color represents mean *CIITA* CpG methylation. **H.** HLA II score and CIITA methylation for pre-B-ALL and T-ALL subtypes is shown for Hemap GSE49031 dataset. Dot color represents methylation of the cg04945379 *CIITA* probe. **I.** Representative flow cytometry plots of MOLM13 cells after 72 h treatment with DMSO or 100 nM decitabine (DAC) and 10 ng/mL IFNγ.

Figure S5

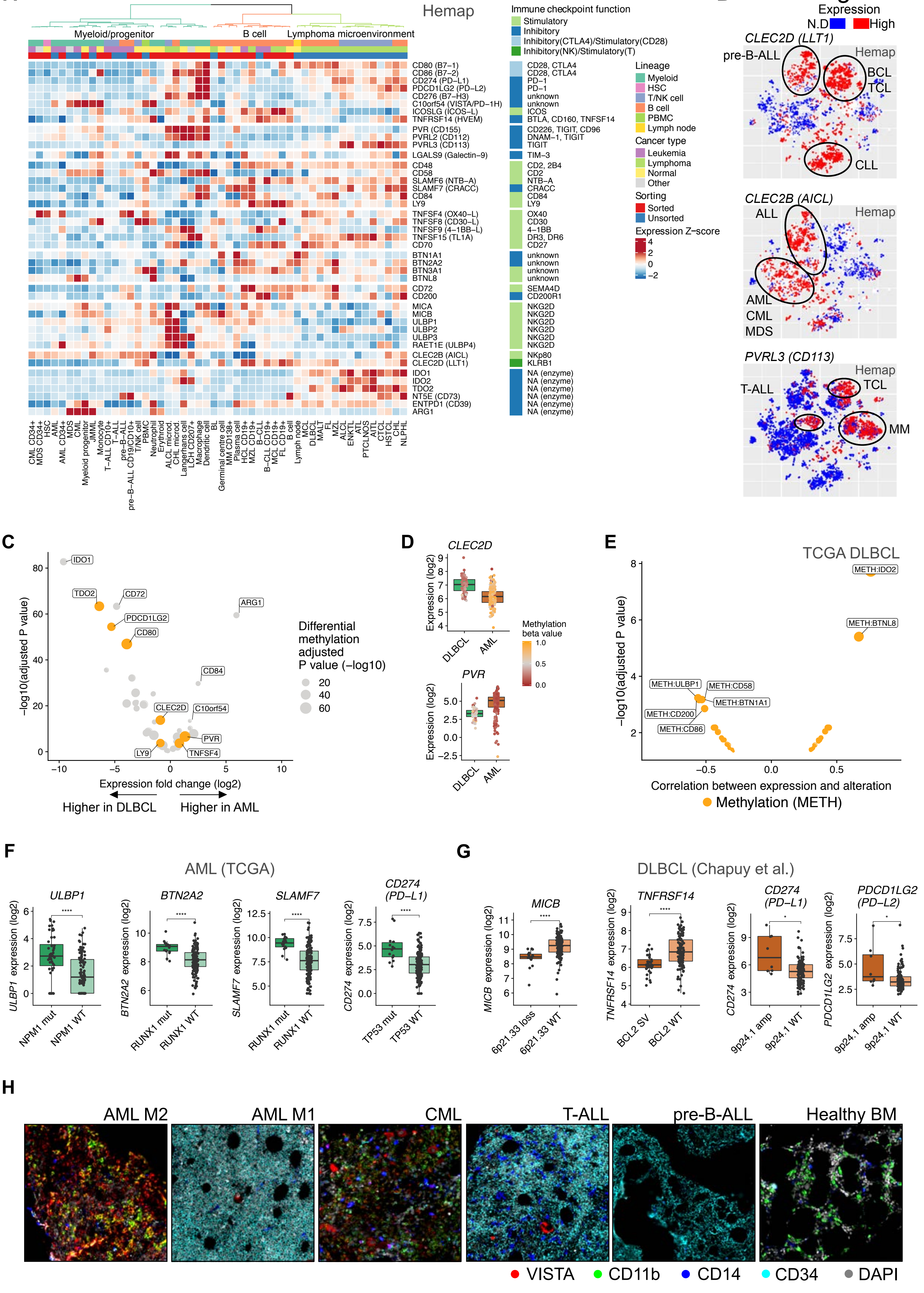
A
Hemap
Myeloid/progenitor
B cell
Lymphoma microenvironment
Immune checkpoint function
Stimulatory
Inhibitory
Inhibitory(CTLA4)/Stimulatory(CD28)
Inhibitory(NK)/Stimulatory(T)
CD80 (B7−1)
CD86 (B7−2)
CD274 (PD−L1)
PDCD1LG2 (PD−L2)
CD276 (B7−H3)
C10orf54 (VISTA/PD−1H)
ICOSLG (ICOS−L)
TNFRSF14 (HVEM)
PVR (CD155)
PVRL2 (CD112)
PVRL3 (CD113)
LGALS9 (Galectin−9)
CD48
CD58
SLAMF6 (NTB−A)
SLAMF7 (CRACC)
CD84
LY9
TNFSF4 (OX40−L)
TNFSF8 (CD30−L)
TNFSF9 (4−1BB−L)
TNFSF15 (TL1A)
CD70
BTN1A1
BTN2A2
BTN3A1
BTNL8
CD72
CD200
MICA
MICB
ULBP1
ULBP2
ULBP3
RAET1E (ULBP4)
CLEC2B (AICL)
CLEC2D (LLT1)
IDO1
IDO2
TDO2
NT5E (CD73)
ENTPD1 (CD39)
ARG1
CD28, CTLA4
CD28, CTLA4
PD−1
PD−1
unknown
unknown
ICOS
BTLA, CD160, TNFSF14
CD226, TIGIT, CD96
DNAM−1, TIGIT
TIGIT
TIM−3
CD2, 2B4
CD2
NTB−A
CRACC
CD84
LY9
OX40
CD30
4−1BB
DR3, DR6
CD27
unknown
unknown
unknown
unknown
SEMA4D
CD200R1
NKG2D
NKG2D
NKG2D
NKG2D
NKG2D
NKG2D
NKp80
KLRB1
NA (enzyme)
NA (enzyme)
NA (enzyme)
NA (enzyme)
NA (enzyme)
NA (enzyme)
Lineage
Myeloid
HSC
T/NK cell
B cell
PBMC
Lymph node
Cancer type
Leukemia
Lymphoma
Normal
Other
Sorting
Sorted
Unsorted
Expression Z−score
4
2
0
−2
CML CD34+
MDS CD34+
HSC
AML
AML CD34+
MDS
CML
Myeloid progenitor
JMML
Monocyte
T−ALL CD10+
T−ALL
pre−B−ALL
pre−B−ALL CD19/CD10+
T/NK cell
PBMC
Neutrophil
Erythroid
ALCL microd.
CHL microd.
Langerhans cell
LCH CD207+
Macrophage
Dendritic cell
BL
Germinal centre cell
MM CD138+
Plasma cell
HCL CD19+
MZL CD19+
B−CLL
B−CLL CD19+
MCL CD19+
FL CD19+
B cell
Lymph node
MCL
DLBCL
MALT
FL
MZL
ALCL
ENKTL
ATL
PTCLNOS
AITL
CTCL
HSTCL
CHL
NLPHL
B
Expression
N.D
High
CLEC2D (LLT1)
pre-B-ALL
Hemap
BCL
TCL
CLL
CLEC2B (AICL)
ALL
Hemap
AML
CML
MDS
PVRL3 (CD113)
Hemap
T-ALL
TCL
MM
C
IDO1
TDO2
CD72
ARG1
PDCD1LG2
CD80
CD84
CLEC2D
C10orf54
PVR
LY9
TNFSF4
−log10(adjusted P value)
80
60
40
20
0
−10
−5
0
5
10
Expression fold change (log2)
Higher in DLBCL
Higher in AML
Differential methylation adjusted P value (−log10)
20
40
60
D
CLEC2D
Expression (log2)
DLBCL
AML
PVR
Expression (log2)
DLBCL
AML
Methylation beta value
1.0
0.5
0.0
E
TCGA DLBCL
METH:IDO2
METH:BTNL8
METH:ULBP1
METH:CD58
METH:BTN1A1
METH:CD200
METH:CD86
−log10(adjusted P value)
Correlation between expression and alteration
Methylation (METH)
F
AML (TCGA)
ULBP1
ULBP1 expression (log2)
NPM1 mut
NPM1 WT
BTN2A2
BTN2A2 expression (log2)
RUNX1 mut
RUNX1 WT
SLAMF7
SLAMF7 expression (log2)
RUNX1 mut
RUNX1 WT
CD274 (PD−L1)
CD274 expression (log2)
TP53 mut
TP53 WT
G
DLBCL (Chapuy et al.)
MICB
MICB expression (log2)
6p21.33 loss
6p21.33 WT
TNFRSF14
TNFRSF14 expression (log2)
BCL2 SV
BCL2 WT
CD274 (PD−L1)
CD274 expression (log2)
9p24.1 amp
9p24.1 WT
PDCD1LG2 (PD−L2)
PDCD1LG2 expression (log2)
9p24.1 amp
9p24.1 WT
H
AML M2
AML M1
CML
T-ALL
pre-B-ALL
Healthy BM
VISTA
CD11b
CD14
CD34
DAPI

**Figure S5. Genetic and epigenetic alterations and cancer subtypes associated with expression of immunomodulatory genes. Related to Figure 5. A.** Expression levels (Z-scores of median log2 expression) of co-stimulatory, co-inhibitory, and other immunomodulatory genes across Hemap cancer samples and normal cell types shown as a heatmap. Rows and columns are clustered using Spearman correlation distance and Ward's linkage. Corresponding receptors and the nature of the interaction (stimulatory/inhibitory) is shown according to Table S5. **B.** Expression of selected cancer type-specific immunomodulatory genes colored on Hemap t-SNE map. Circles indicate clusters containing cancer types expressing the indicated gene. N.D = Not detected. **C.** Volcano plot of differentially expressed immunomodulatory genes between TCGA AML and DLBCL. Point size indicates negative log10 of *P* value of differential methylation of the gene between the two cancer types. Genes with methylation log2 fold change > 1.5 are colored in yellow. **D.** Comparison of expression of *CLEC2D* and *PVR* between TCGA DLBCL and TCGA AML. Dots indicating individual patients are colored by average methylation within 1 kb of the transcription start site. **E.** Volcano plot of correlations (Spearman) of immunomodulatory gene expression with promoter methylation in TCGA DLBCL. Point size is proportional to the adjusted *P* value. **F.** Selected immunomodulatory genes whose expression is correlated with genetic alterations in TCGA AML are shown as boxplots comparing cases with alteration (mut) and wild-type (WT) cases. Nominal *P* values obtained using two-sided Wilcoxon rank sum test are shown. **G.** Selected immunomodulatory genes whose expression is correlated with genetic alterations in DLBCL (GSE98588) are shown as boxplots comparing cases with alteration (mut/SV/amp) and wild-type (WT) cases. Nominal *P* values obtained using two-sided Wilcoxon rank sum test are shown. **H.** Multiplex immunohistochemistry of BM biopsies for VISTA, CD11b, CD14, CD34 and DAPI from representative patients of each profiled cancer type and healthy controls.

### Figure S6

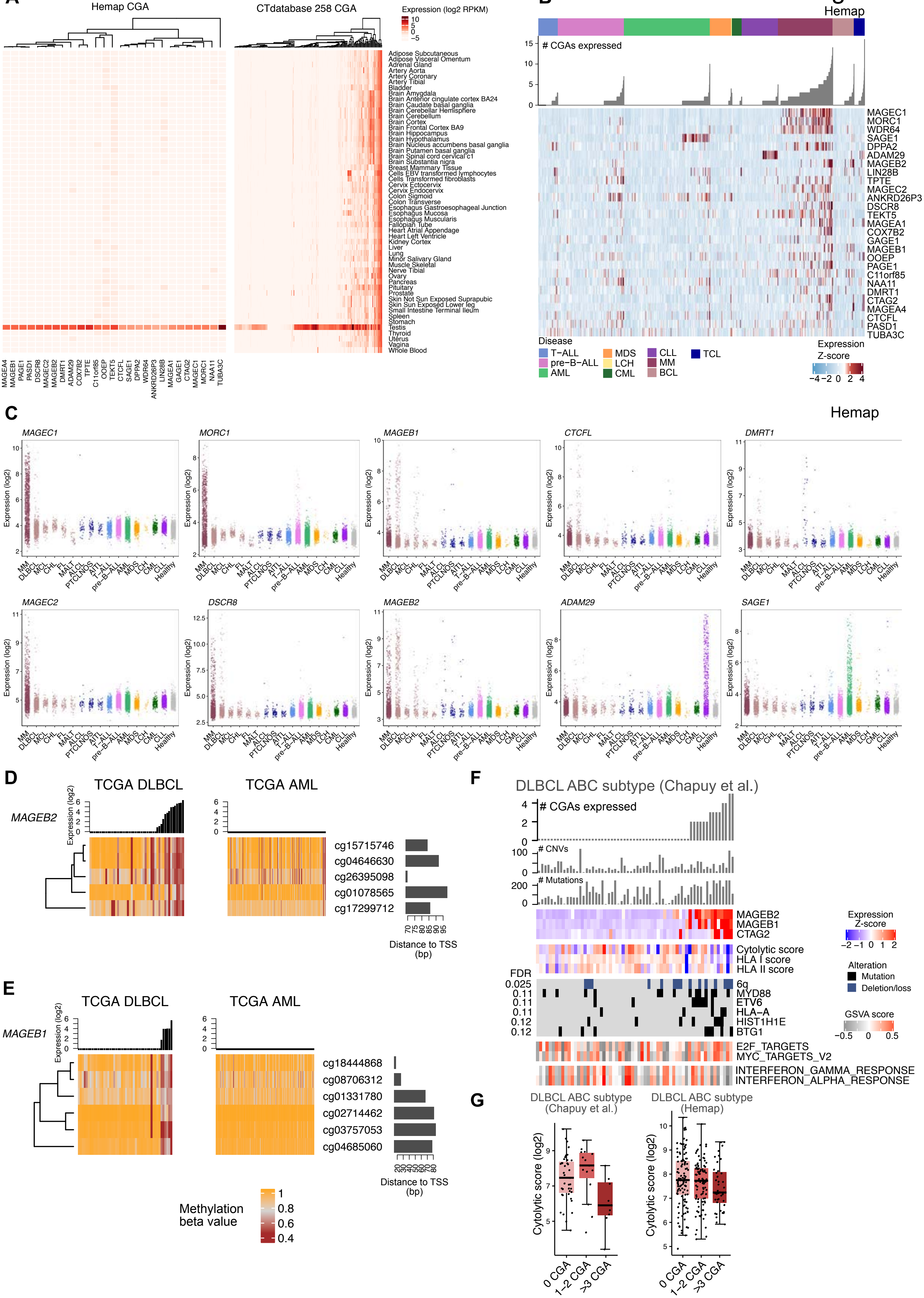
A
Hemap CGA
GTEx
CTdatabase 258 CGA
Expression (log2 RPKM)
B
Hemap
### CGAs expressed
Disease
T−ALL
pre−B−ALL
AML
MDS
LCH
CML
CLL
MM
BCL
TCL
Expression
Z-score
C
Hemap
MAGEC1
MORC1
MAGEB1
CTCFL
DMRT1
MAGEC2
DSCR8
MAGEB2
ADAM29
SAGE1
Expression (log2)
D
TCGA DLBCL
TCGA AML
MAGEB2
Distance to TSS (bp)
E
MAGEB1
Methylation beta value
F
DLBCL ABC subtype (Chapuy et al.)
### CGAs expressed
### CNVs
### Mutations
Cytolytic score
HLA I score
HLA II score
FDR
Alteration
Mutation
Deletion/loss
GSVA score
E2F_TARGETS
MYC_TARGETS_V2
INTERFERON_GAMMA_RESPONSE
INTERFERON_ALPHA_RESPONSE
G
DLBCL ABC subtype (Chapuy et al.)
DLBCL ABC subtype (Hemap)
Cytolytic score (log2)
0 CGA
1−2 CGA
>3 CGA

**Figure S6. CGA expression in hematological malignancies. Related to Figure 6. A.** GTEx dataset log2 RPKM expression of 27 CGAs found expressed in hematological malignancies are shown as a heatmap. On the right expression of CTdatabase CGA genes (total 258) are shown. **B.** Expression of CGAs across hematological malignancies in Hemap are shown as a heatmap. Total number of expressed CGAs per patient are shown as barplots. **C.** Log2 expression for selected CGAs for Hemap cancers and healthy controls. **D-E.** MAGEB2 and MAGEB1 methylation values are shown for each probe associated with the gene for TCGA DLBCL and AML data sets. Samples are ranked by gene expression indicated above the heatmap. Probe distances (bp) from TSS are shown as barplot on the right. **F.** Genetic alterations and gene sets correlated with the number of expressed CGAs in the ABC subtype of DLBCL (Chapuy *et al.*). Mutation and CNV load, immunological scores, and expression of 3 most commonly expressed CGAs in DLBCL are shown. **G.** Cytolytic score stratified by CGA number in ABC DLBCL (Chapuy *et al.* and Hemap).

Figure S7

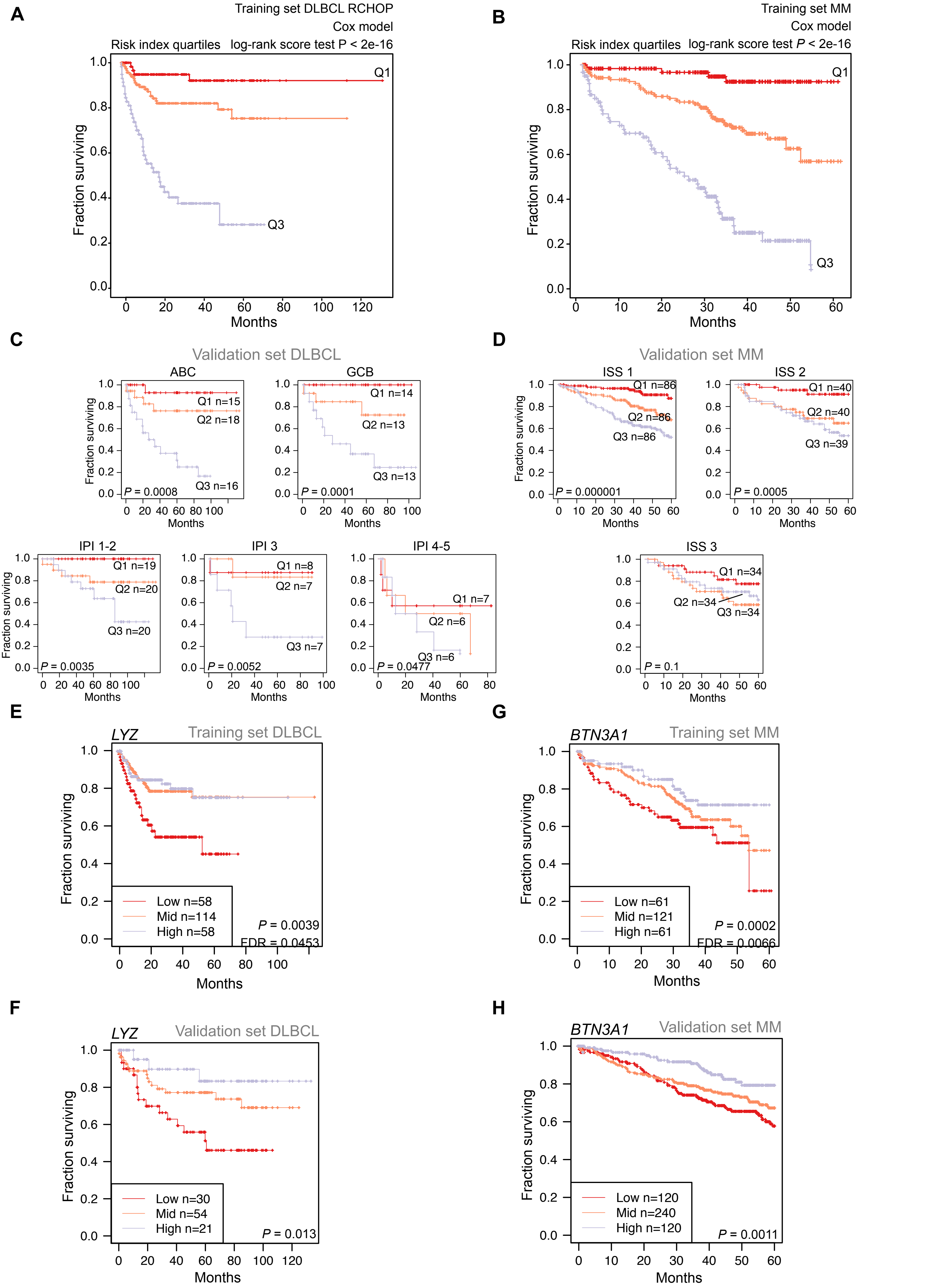

A
Training set DLBCL RCHOP
Cox model
Risk index quartiles log-rank score test P < 2e-16
Q1
Q3
Fraction surviving
Months
B
Training set MM
Cox model
Risk index quartiles log-rank score test P < 2e-16
Q1
Q3
Fraction surviving
Months
C
Validation set DLBCL
ABC
Q1 n=15
Q2 n=18
Q3 n=16
P = 0.0008
GCB
Q1 n=14
Q2 n=13
Q3 n=13
P = 0.0001
IPI 1-2
Q1 n=19
Q2 n=20
Q3 n=20
P = 0.0035
IPI 3
Q1 n=8
Q2 n=7
Q3 n=7
P = 0.0052
IPI 4-5
Q1 n=7
Q2 n=6
Q3 n=6
P = 0.0477
Fraction surviving
Months
D
Validation set MM
ISS 1
Q1 n=86
Q2 n=86
Q3 n=86
P = 0.000001
ISS 2
Q1 n=40
Q2 n=40
Q3 n=39
P = 0.0005
ISS 3
Q1 n=34
Q2 n=34
Q3 n=34
P = 0.1
Fraction surviving
Months
E
LYZ
Training set DLBCL
Low n=58
Mid n=114
High n=58
P = 0.0039
FDR = 0.0453
Fraction surviving
Months
G
BTN3A1
Training set MM
Low n=61
Mid n=121
High n=61
P = 0.0002
FDR = 0.0066
Fraction surviving
Months
F
LYZ
Validation set DLBCL
Low n=30
Mid n=54
High n=21
P = 0.013
Fraction surviving
Months
H
BTN3A1
Validation set MM
Low n=120
Mid n=240
High n=120
P = 0.0011
Fraction surviving
Months

**Figure S7. Survival associations of immunological features. Related to Figure 7. A.** Kaplan-Meier curves of overall survival for DLBCL patients stratified by the risk index of the immunological risk model in the training set (Hemap R-CHOP treated GSE10846,GSE17372). Patients were divided into three groups based on quartiles and *P* value was obtained using the score (log-rank) test. **B.** Kaplan-Meier curves of overall survival for MM patients stratified by the risk index of the immunological risk model in the training set (GSE19784). Patients were divided into three groups based on quartiles and *P* value was obtained using the score (log-rank) test.**C.** Kaplan-Meier curves of overall survival for DLBCL patients stratified by the risk index of the immunological risk model in the validation set (GSE98588), divided into subgroups based on existing prognostic markers, ABC/GCB subtype and IPI. Patients were divided into three equally-sized groups and *P* value was obtained using the score (log-rank) test. **D.** Kaplan-Meier curves of overall survival for MM patients stratified by the risk index of the immunological risk model in the validation set (GSE16716 and GSE24080), divided into subgroups based on the existing prognostic marker ISS. Patients were divided into three equally-sized groups and *P* value was obtained using the score (log-rank) test. **E.** Kaplan-Meier curves of overall survival for DLBCL patients stratified by *LYZ* expression in the training set (GSE10846, GSE17372). Patients were divided into three groups based on quartiles and *P* value was obtained using the Wald test. **F.** Kaplan-Meier curves of overall survival for DLBCL patients stratified by *LYZ* expression in the validation set (GSE98588). Patients were divided into three groups based on quartiles and *P* value was obtained using the Wald test. **G.** Kaplan-Meier curves of overall survival for MM patients stratified by *BTN3A1* expression in the training set (GSE19784). Patients were divided into three groups based on quartiles and *P* value was obtained using the Wald test. **H.** Kaplan-Meier curves of overall survival for MM patients stratified by *BTN3A1* expression in the validation set (GSE16716 and GSE24080). Patients were divided into three groups based on quartiles and *P* value was obtained using the Wald test.

#### SUPPLEMENTAL TABLES

**Table S1. Sample annotation table and immunological scores of Hemap samples included in the study. Related to Figure 1.**

(A) Sample annotation table and immunological scores of Hemap samples
(B) Flow cytometry panel for cytolytic score validation

**Table S2. Associations of cytolytic score to gene expression and mIHC validation. Related to Figure 2.**

(A) Genes correlated to cytolytic score in hematological malignancies
(B) Multiplexed immunohistochemistry antibody panel
(C) Multiplexed immunohistochemistry data

**Table S3. Genomic and clinical correlations of cytolytic score. Related to Figure 3.**

(A) TCGA AML genomic and clinical correlations
(B) BeatAML genomic and clinical correlations
(C) Chapuy *et al*. DLBCL genomic and clinical correlations
(D) Chapuy *et al.* DLBCL ABC genomic and clinical correlations
(E) Chapuy *et al.* DLBCL GCB genomic and clinical correlations

**Table S4. Genomic and clinical correlations of HLA expression. Related to Figure 4.**

(A) HLA II score development. Differentially expressed genes between antigen-presenting cells (APC) and other normal hematopoietic cells (non-APC).
(B) Flow cytometry panels for HLA II score validation
(C) TCGA AML genomic and clinical correlations with HLA scores
(D) TCGA AML CIITA genomic and clinical correlations
(E) BeatAML genomic and clinical correlations with HLA scores

**Table S5. Genomic, clinical, and cancer type correlations of immune checkpoint gene expression. Related to Figure 5.**

(A) List of co-stimulatory and co-inhibitory immune checkpoint genes and other immunomodulators
(B) Hemap cancer subtype associations
(C) Differential expression and methylation between TCGA AML and DLBCL
(D) TCGA AML genomic and clinical correlations
(E) BeatAML genomic and clinical correlations

(F) TCGA DLBCL genomic and clinical correlations

(G) Chapuy *et al.* DLBCL genomic and clinical correlations

(H) Multiplexed immunohistochemistry antibody panel

(I) Multiplexed immunohistochemistry data

**Table S6. Genomic and clinical correlations of CGA expression. Related to Figure 6.**

(A) Hemap MM GSEA

(B) Chapuy et al. DLBCL genomic and clinical correlations

(C) Chapuy et al. DLBCL ABC genomic and clinical correlations

(D) Chapuy et al. DLBCL GCB genomic and clinical correlations

(E) Chapuy et al. DLBCL ABC GSEA

(F) Chapuy et al. DLBCL GCB GSEA

**Table S7. Survival analyses of immunological features. Related to Figure 7.**

(A) Univariate survival associations in DLBCL training data (Hemap).

(B) Univariate survival associations in MM training data (Hemap).

(C) Univariate survival associations in Hemap AML.

(D) Univariate survival associations in TCGA AML.
